# Expanding the MR1 ligandome using chemical class-specific fragmentation and molecular networking

**DOI:** 10.1101/2025.05.29.656472

**Authors:** Chiara Lecchi, Alessandro Vacchini, Stefano Sainas, Marco L. Lolli, Matthew Luedtke, Lucia Mori, Gennaro De Libero, Silvia Balbo, Peter W Villalta

**Author notes:** **Corresponding Authors Peter W Villalta** - Masonic Cancer Center and Department of Medicinal Chemistry, University of Minnesota; Minneapolis, MN 55455, United States, Address reprint requests to Peter Villalta, 2231 6^th^ Ave SE. 2-143 CCRB, **Gennaro De Libero** - Experimental Immunology, Department of Biomedicine, University Hospital and University of Basel, Basel 4031, Switzerland, **Silvia Balbo** - Masonic Cancer Center and Division of Environmental Health Sciences, School of Public Health, University of Minnesota; Minneapolis, MN 55455, United States. The authors contributed equally to the work presented (Chiara Lecchi, Alessandro Vacchini).

## Abstract

The identification and subsequent characterization of unknown analytes using mass spectrometry presents a long-standing challenge across many research fields, particularly when analyte levels are low and the compound class is underrepresented in mass spectral databases. We have developed a data analysis workflow for investigating classes of small molecules and demonstrated its application through the reanalysis of data collected to probe for modified nucleoside MR1-presented antigens. We reanalyzed the datasets to screen for additional classes of compounds within the MR1 ligandome using Compound Discoverer, a commercial software package designed for metabolomic analysis, featuring fragmentation filtering nodes, molecular networking, and spectral database searching. Our study identified two compound classes that bind to MR1. One class includes compounds characterized by the presence of a ribityl substructure and molecular formulas consistent with structural similarity to riboflavin, where the most abundant compound differs from riboflavin by two additional oxygen atoms and one fewer carbon atom. A second class comprises an adenosine monophosphate isomer and larger analytes that are putatively identified as consisting of di- and tri-covalently bound nucleotides. The application of our analytical approach to characterize the MR1 ligandome demonstrates the power of combining compound-class fragmentation, molecular networking, and mass spectral database searching in exploring receptor ligandomes and, more generally, identifying novel classes of compounds.

## INTRODUCTION

T lymphocytes target cells by recognizing antigens bound to cell-surface proteins encoded by the major histocompatibility complex (MHC). Immunopeptidomics is an LC-MS-based method for identifying peptide antigens that bind to classical MHC proteins.^1, 2^ A nonclassical MHC class I-like molecule, called MHC class I-related protein 1 (MR1), presents small molecule ligands as antigens to a population of T cells, including mucosal-associated invariant T (MAIT) cells. However, the nature of these ligands remains largely underexplored and requires analytical approaches capable of identifying trace levels of unknown small molecules presented by MR1. The identification and subsequent characterization of unknown small molecules using mass spectrometry is a long-standing challenge shared by many research fields, including natural product screening,^3^ exposomics,^4^ and metabolomics.^5^ The difficulty increases when the analytes are at trace levels and not included in fragmentation databases. Trace-level detection of analytes can be significantly facilitated by leveraging the well-established inverse relationship between flow rate and electrospray ionization and sampling efficiency,^6^ as seen in immunopeptidomics.^7^ This is not typically done in standard small-molecule screening assays; however, it has been implemented in our DNA adductomics methodology^8-10^ to study DNA damage by screening for covalently modified nucleosides (DNA adducts) detected in hydrolyzed DNA samples. Additionally, compound-class diagnostic fragmentation patterns are utilized to screen and characterize the covalently modified nucleosides and nucleobases by taking advantage of the near-universal neutral loss of the 2’-deoxyribose and common neutral losses of nucleobases, along with the appearance of one of the protonated nucleobases.^11^

Our DNA adductomics method has previously been modified to probe for modified nucleoside ligands captured by MR1,^12^ and several specific carbonyl-nucleobase adducts were ultimately identified^12^. Other significant efforts have been made in the past decade to describe MR1 ligands, including seminal work indicating that microbial riboflavin metabolism is a source of MR1 ligands that activate mucosal-associated invariant T (MAIT) cells.^13, 14^ Results from this line of work have primarily identified ribityllumazines and ribityl-uracil derivatives as the prominent representatives of MAIT cell-stimulatory ligands.^15, 16^ The extent of the known MR1 ligandome has recently been reviewed,^17^ including the recent identification of vitamin B6-related compounds.^18^

In the work presented here, we reanalyzed our MR1 study nanoLC-MS/MS data^12^ using compound class-specific fragmentations in conjunction with molecular networking^19, 20^ and spectral database searching to identify and characterize additional MR1-binding molecules. For this reanalysis, we anticipated class-specific diagnostic fragmentation for ribityl-containing compounds to identify new ribityl-containing species that preferentially bind to MR1 and interrogated the data for nucleobase neutral losses and product ions to identify a novel class of MR1-interacting molecules consisting of ribose-nucleotides. This study is meant to serve as a proof-of-principle for the ability of our method to elucidate the ligandome of mammalian cell receptors.

## MATERIALS AND METHODS

### Chemicals and Supplies

Acetonitrile (ACN, LC-MS grade), water (LC-MS grade), and formic acid (FA, 98% v/v) were purchased from Fluka (St. Louis, MO, USA). Microcon single-membrane filtration devices (10- kDa cutoff, 0.5 mL) were purchased from Millipore-Sigma (Billerica, MA, USA). Silanized vials were purchased from ChromTech (Apple Valley, MN, USA). Di-adenine RNA (ApAp) was purchased from Horizon Discovery Biosciences Limited (Cambridge, UK). All other chemicals and supplies were purchased from Sigma-Aldrich or Fisher Scientific. Chemical reagents for synthesis were obtained from commercial sources (Sigma Aldrich and BLD Pharma) and used without further purification. Analytical-grade solvents (diethyl ether, dichloromethane [DCM], methanol [MeOH], tert-butyl alcohol [*t*-ButOH]) were also used without further purification. When necessary, the solvents were dried over 4 Å molecular sieves. Anhydrous MeOH was obtained by distillation over CaCl_2_ and stored over activated 3 A molecular sieves for a minimum of 24 h. Anhydrous pyridine was obtained by distillation over CaH_2_. Thin layer chromatography (TLC) was performed on silica gel-coated 5 × 20 cm plates at a layer thickness of 0.25 mm, visualized using UV light (254 nm) or TLC visualization reagents (KMnO_4_, ninhydrin, iodine, and 2,4- dinitrophenylhydrazine). Anhydrous Na_2_SO_4_ was utilized as a drying agent for the organic phases.

### Synthesis of P^1^P^2^-diadenosine-5’-pyrophosphate (AppA)

The synthesis of AppA was conducted using a method previously reported in the literature^21^ and illustrated in Scheme S1. Details of the synthesis and the compound’s NMR characterization (Figures S1 and S2) are included in the Supporting Information.

### Production of soluble MR1 and sCD1d proteins

Analyzed samples were obtained by eluting molecules bound to recombinant soluble human CD1d (sCD1d) and K43A-MR1 (sMR1) expressed in CHO-K1 cells.^12, 22^ Proteins were produced by culturing cells in serum-free medium (MAM-PF2, Cat#10-02F25-I) supplemented with 1 mM sodium pyruvate (Cat#5-60F00-H), 2 mM stable glutamine (Ala-Gln, Cat#5-10K50-H), 1% non-essential amino acids (NEAA, Cat#5-13K00-H), and 50 µg/mL kanamycin (Cat#4-08F00-H) (all from Bioconcept). The collected supernatants were centrifuged for 5 minutes at 600g and then filtered through a 0.22 µm filter. Recombinant proteins were quantified by ELISA as previously described.^12^

### Elution of sMR1and sCD1d-bound compounds and sample preparation

sMR1 and sCD1d proteins were purified with HiTrap Fibro™ PrismA (Cytiva, Cat#17549855) by washing with 20 column volumes (CV) of PBS, then with 10 CV of mass-spectrometry grade water adjusted to pH 7.2 with ammonium carbonate (Sigma-Aldrich, Cat#11204). A solution of 20% methanol and 0.1% formic acid was used to elute the ligands from the column-bound sMR1 and sCD1d. The eluted material was pooled in silanized glass vials (Thermo Fisher Scientific, Cat#SAA-SV2-2) and the volume was reduced under N_2_ stream, followed by a previously described clean-up procedure.^23^ Briefly, 50 μL of concentrated material eluted from sMR1 and sCD1d were pre-chilled in 2 mL Eppendorf tubes on ice. Three volumes (150 μL) of cold acetonitrile:acetone (1:1, v/v) were added to the samples, and after being vortexed for 1 minute, they were left on ice for 10 minutes. Then, the samples were centrifuged at 16,000g for 10 minutes at 4°C, and the supernatants were transferred to new silanized glass vials. The samples were finally dried in a SpeedVac centrifuge and reconstituted in 50 μL LC-MS water, and 5 μL were injected into the LC-MS system.

### Cell medium preparation for LC-MS analysis

Cell medium was used as a negative control and prepared as follows. 100 μL of cell medium and 400 μL of water were mixed and added to a 1 mL 30 mg Strata-X (Phenomenex, Torrance, CA) solid-phase extraction cartridge, which had been preconditioned with three 1 mL aliquots of methanol followed by 1 mL water. The cartridge was washed with three 1 mL aliquots of water, followed by elution of the analytes with two 1 mL aliquots of methanol containing 0.1% formic acid. The collected volume was then dried in a SpeedVac. Before LC-MS analysis, the samples were reconstituted in 100 μL of LC-MS water. Five μL were injected into the system.

### LC conditions for LC-MS analysis

All LC-MS analyses of purified samples were conducted on a Thermo Orbitrap Lumos mass spectrometer coupled with a Dionex RSLCnano HPLC featuring a Nanospray Flex ion source (Thermo Scientific, Waltham, MA). A capillary column (200 mm x 75 μm, New Objective, Woburn, MA) was hand-packed using reversed-phase Luna C18 (5 μm, 120 Å, Thermo Scientific, Waltham, MA), and separation was achieved at room temperature with a flow rate of 0.3 μL/min, employing 0.05% formic acid as mobile phase A and acetonitrile as mobile phase B. Chromatographic separation utilized gradient elution, starting with 2% B for the initial 5.5 min at a flow rate of 1 μL/min. The flow rate was decreased to 0.3 μL/min over 0.5 min, with the injection valve position switching at 6 min into the run to remove the 5 μL injection loop from the flow path, followed by a linear gradient increasing from 2% to 50% B over 39 min, culminating in a rise to 98% B over 1 min and a 2 min hold. The electrospray voltage was set at 2.2 kV, with a source temperature of 350°C and an S-Lens RF setting of 50%.

### MS parameters for Data Dependent-Constant Neutral Loss/MS^3^ (DDA-CNL/MS^3^) analysis

A high-resolution data-dependent MS^3^ neutral loss/product ion screening approach was employed for this analysis. This branched method involves monitoring neutral losses and the emergence of specific ions listed in Table S1, activated by the observation of one of them through MS3 fragmentation. For the MS^1^ full scans, the Orbitrap analyzer operated within a mass range of 150-1000 Da, a resolution setting of 120000 (at *m/z* 200), an automatic gain control (AGC) setting of 1 × 10^6^ (250%), and a maximum injection time of 50 ms. A stepped MS^2^ HCD (25, 50%) fragmentation was executed using a “Top Speed” setting of 3 s with quadrupole isolation (1.5 Da), Orbitrap detection at a resolution of 15000, a maximum injection time of 50 ms, an AGC target of 2 × 10^5^ (400%), and a minimum intensity threshold of 20000. Dynamic exclusion was implemented with an exclusion duration of 15 s and a mass tolerance of ± 5 ppm. MS^2^ fragment ions resulting from the characteristic neutral losses or the production of specific fragment ions cited in Table S1 (± 5 ppm) underwent MS^3^ HCD (30%) fragmentation and Orbitrap detection at a resolution of 15000, with MS^1^ quadrupole isolation of 1.5 Da, MS^2^ ion trap isolation of 2 Da, a maximum injection time of 200 ms, and an AGC target of 200000 (400%).

### MS parameters for DDA-CNL/MS^3^ at high HCD with inclusion list

The analysis, which included an inclusion list and a high HCD value, was performed using the DDA-CNL/MS^3^ parameters reported above, with the following exceptions. An inclusion list of 56 *m/z* values (Table S2) was used, consisting of the *m/z* values found using the screening DDA- CNL/MS^3^ analysis (Table S3), ^13^C_4_,^15^N_2_-riboflavin, and positive ion *m/z* values corresponding to previously observed ribityl-containing analytes.^24^ The only MS^3^ triggering criterion was the neutral loss of 134.0579 (ribityl) with a 25 ppm mass tolerance. The MS^2^ and MS^3^ collision energies were set to 55% HCD.

### Data Analysis for Analyte Screening with Molecular Networking

Analyte screening was conducted using Compound Discoverer (CD) software (Ver. 3.3.3.200, Thermo Scientific, Waltham, MA), with the workflow graphically illustrated in Scheme S2 and a complete set of parameters detailed in the Supporting Information. The following are the most significant parameters. The “Input Files” node was used to submit spectra from the entire chromatographic run to the “Select Spectra” node, followed by the “Detect Compounds” node with the following parameters: Mass Tolerance, 5 ppm; Min. Peak Intensity, 1000; Precursor Mass Tolerance, 0.025 Da; Chromatographic S/N Threshold, 1.5; Remove Baseline, False; Ions, [M+H] +1. The “Compound Class Scoring” node utilized the targeted masses listed in Table S1 with the following parameters: Mass Tolerance, 5 ppm; RT Tolerance, 0.5 min.; Minimum Valley, 10%. The “Predict Compositions” node had the following parameters: Mass Tolerance, 5 ppm; Max. Element Counts, C90 H190 Br Cl Fe N10 O18 P3 S5; Max. RDBE, 40; Max. H/C 3.5. The “Search Neutral Losses” node utilized a neutral loss mass of 134.05791 Da (C_5_H_10_O_4_), and the “Compound Class Scoring” node used the product ion of *m/z* 136.06177 (adenine, C_5_H_5_N_5_·H^+^). This node had the following parameters: high accurate mass tolerance of 5 ppm and low accurate mass tolerance of 20 ppm. The “Assign Compound Annotations” node maintained a 5 ppm mass tolerance. The “Search mzVault” and “Search mzCloud” nodes applied mass tolerances of 10 ppm. The “Search mzVault” node searched the NIST 2020 MSMS High Resolution and EPA Comptox NIST Highres 20210802 databases. The post-processing “Differential Analysis” node was employed to generate ratios corresponding to the signal intensities of the analytes in the sMR1 and sCD1d samples. The “Generate Molecular Networks” node created molecular networks between analytes observed in the CD workflow. The resulting “Compounds” identification output from the CD data processing workflow was filtered using the “Result Filters” feature to identify analytes more abundant in the treatment samples by filtering the “Compounds” with “AND” logic operator to combine “Neutral Losses” (is true, in any loss) and “Ratio” (is greater than 5), where the value represents the ratio of the “Area”s of the sMR1 samples over the sCD1d samples. The molecular network (MN) feature was used to assess structural similarities of the detected analytes. The following parameters were adjusted to fine-tune and investigate the MN output, with the values used in the final summaries listed in the figure captions: 1) the “Score” parameter is the minimum MS^n^ similarity score value necessary for a connection to a similar compound; 2) the “Coverage” parameter is the minimum “Forward or Reverse coverage” value for a given connection; 3) the “Matched Fragments” value specifies the minimum number of fragment ions that match between two connected features.

## RESULTS AND DISCUSSION

### Identification of the Ribityl-Containing Compounds

The report of ribityl-containing compounds binding to MR1 in bacterial systems^24^ led us to examine the previously collected mammalian MR1 LC-CNL/MS^3^ data^12^ for the presence of ribityl- containing compounds. Based on the structural similarities of the ribityl, ribose, and deoxyribose moieties, as well as the fact that deoxyribose is nearly universally lost upon MS^2^ fragmentation of nucleosides,^11^ it was assumed that the neutral loss of the ribityl group would be consistently observed during the fragmentation of ribityl-containing compounds. The CD analysis of the LC- CNL/MS^3^ data for the sMR1 and sCD1d samples identified 7719 features with a minimum peak intensity of 1 × 10^6^, out of which 1138 analytes had a signal intensity sMR1/sCD1d ratio greater than 5. Selecting from the 7719 features for the ribityl moiety neutral loss (134.05791, C_5_H_10_O_4_) through the CD “Search Neutral Losses” node identified 73 analytes. Filtering the 7719 features for the combined ratio and neutral loss criteria resulted in 39 analytes.

### Molecular Networking for Ribityl-Containing Compounds

The CD software’s molecular networking (MN) tool was used to further investigate the detected analytes. The MN map with an sMR1/sCD1d feature intensity ratio greater than 5 is depicted in Figure 1, Panels A, B, and C, with the latter panel showing the MN map after the addition of the ribityl neutral loss criteria. Of the 39 features observed with a ratio greater than 5 and a neutral loss of ribityl, 22 are connected in the MN cluster (Figure 1, Panel C), while the remaining 16 analytes are unconnected due to either low MS^2^ similarity scores (less than 30) and/or low MS^2^ forward/reverse coverage values (less than 50) for MS^2^ fragmentation spectra comparisons. CD as riboflavin through the mzCloud fragmentation database (https://www.mzcloud.org/) search with a ‘Best Match” value of 80.2, suggesting that the other analytes in the cluster are structurally similar to riboflavin. The database spectrum for riboflavin and the experimental spectrum are displayed together in Figure 1, Panel D, in a mirror plot collected using HCD (beam-type) and CID (trap-type) fragmentation, respectively. HCD fragmentation with stepped values of 25% and 50% is more energetic than the CID fragmentation at 25%, likely accounting for the presence of additional product ions In the experimental spectrum.

**Figure 1.**
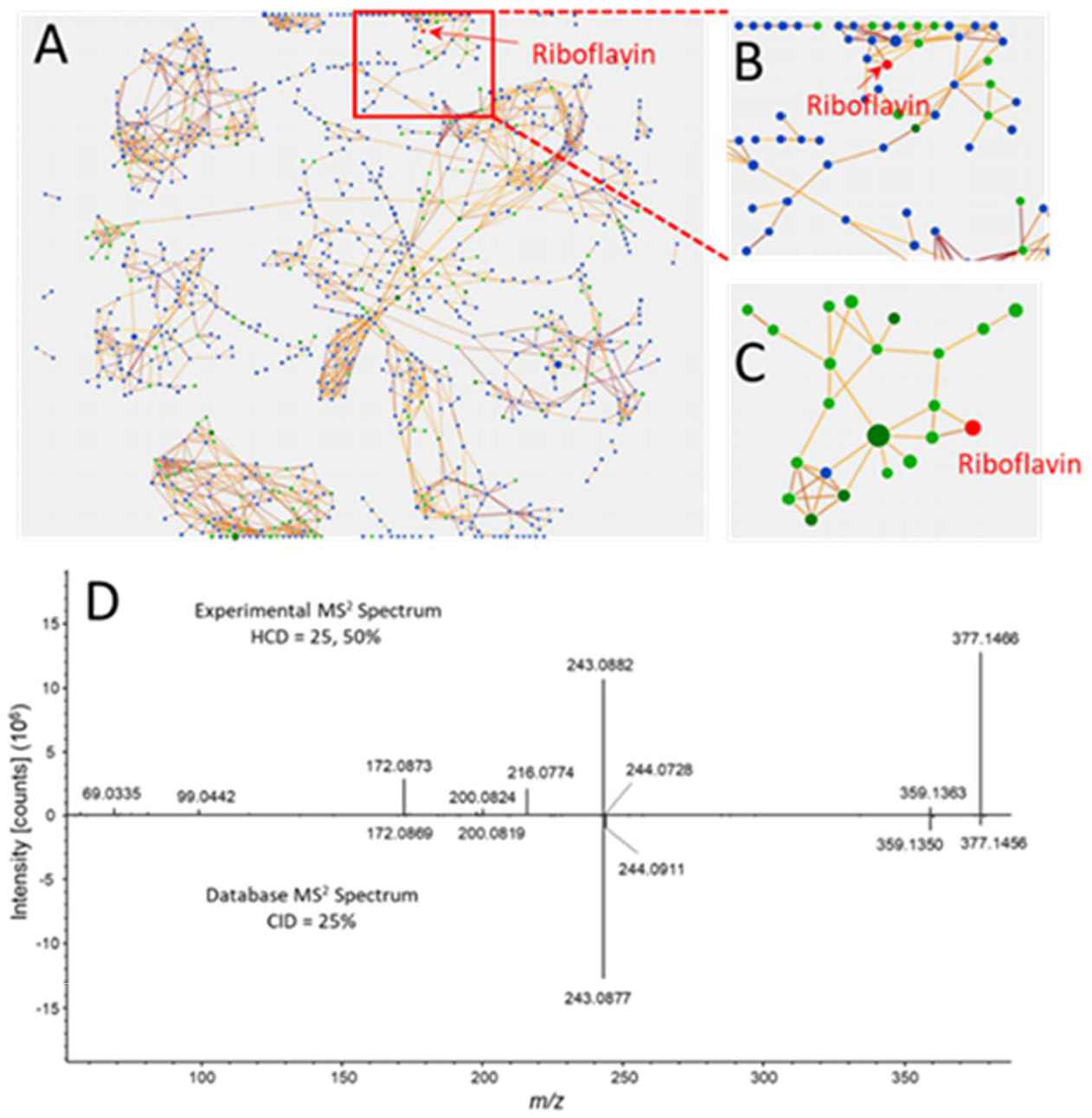
Molecular networking map without orphans shown and thresholds of score=30, coverage=50 and matched fragments=3. Panel A) Full molecular network of analytes with an sMR1/sCD1d ratio>5. Panel B) Expanded region of full molecular network of analytes around the riboflavin node. Panel C) Molecular network with an sMR1/sCD1d ratio>5 and a neutral loss of ribityl. Panel D) Mirror plot of the experimental DDA MS^2^ spectrum of the m/z 377.1461 analyte acquired at a retention time of 23.27 min with stepped (25, 50%) HCD collision energy. Bottom panel: mzCloud database MS^2^ entry for riboflavin collected with a CD fragmentation energy of 25%.

To confirm the identity of this analyte, isotopically labeled ^13^C_4_,^15^N_2_-riboflavin (*m/z* 383.1531) was spiked into the sMR1 and sCD1d samples at a concentration of 1 fmol/μL before reanalyzing them using our DDA-CNL/MS^3^ method with an inclusion list (Table S2) consisting of ^13^C_4_,^15^N_2_-riboflavin, the ribityl-containing analytes identified here, and the analytes previously reported by Hariff et al. in their bacterial sMR1 study.^24^ The MS^2^ data was acquired at higher collision energy (HCD 55) to obtain more extensive fragmentation of the analytes (Supporting Information) and to confirm the identity of riboflavin. The extracted ion chromatograms (Figure 2, Panels A-D) and product ion spectra of the spiked ^13^C_4_,^15^N_2_-riboflavin and the putative riboflavin (Figure 2, Panels E-F) confirm the assignment of the signal to riboflavin and indicate that the level of riboflavin is approximately 10 times higher in the sMR1 sample than in the sCD1d sample. The observation of riboflavin in the mammalian sMR1 samples is consistent with earlier work,^13, 24^ which identified it as the first observed ligand of bacterial sMR1, reinforcing the validity of our experimental setup and the other observations reported in the present study. To further characterize the ribityl-containing features detected, the data were reprocessed using the “Generated Expected Compounds” node of CD with riboflavin as the “Compound Selected” to investigate the possibility that the riboflavin-like analytes are riboflavin metabolites and to assign possible metabolite transformations. Additionally, the Metabolika and BioCyc CD nodes, with the latter using the “human” database, were used to search their corresponding databases for riboflavin- related (ribityl-containing) analyses using the “By Formula or Mass” feature with a mass tolerance of 5 ppm and a maximum of 3 predicted compositions to be searched per compound. Riboflavin was the only ribityl-containing analyte identified here that was attributable to one of the BioCyc or Metabolika pathways.

**Figure 2.**
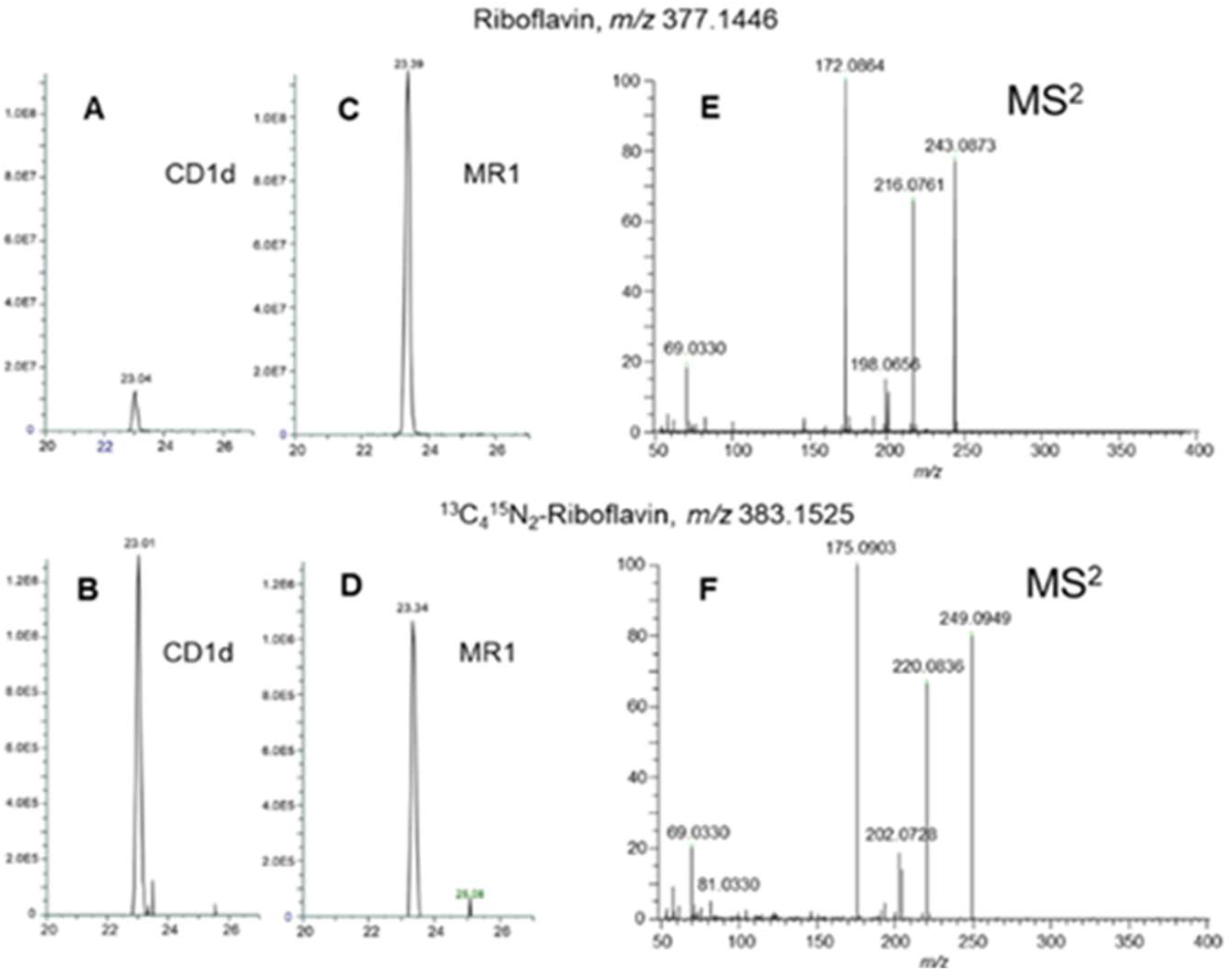
Extracted ion chromatograms and MS^2^ spectra of riboflavin (Panels A.C) and ^13^C_4_,^15^N_2_-riboflavin (Panels B.D) at 5 ppm mass tolerance and their MS^2^ spectra (Panels E,F), respectively, collected at an HCD fragmentation energy of 55%.

The identified ribityl-containing analytes are listed in Table 1, along with their molecular formulas, which are proposed based on the measured molecular masses and assumed structural similarities to riboflavin. Table 1 also includes the differences between the detected masses and those of the proposed molecular formulas, as well as discrepancies in molecular formulas compared to riboflavin. The table caption includes possible metabolic transformation assignments. The chromatographic peak area for each analyte is presented in Table 1 and plotted in Figure S3. The extracted *m/z* ion chromatograms for the analytes in Table 1 were manually examined; their presence in the sMR1 samples and absence in the sCD1d samples were confirmed, except for riboflavin, which was found in a relatively small amount (approximately 10%) in the sCD1d samples, as noted in the CD results. The signal intensity of the 396.1285 Da (*m/z* 397.1348) analyte was ten times higher than that of any other unknown analyte. The peak areas of this analyte and riboflavin (*m/z* 377.1458) accounted for 67% and 14%, respectively, of the total intensities of all the analytes listed in Table 1. The sMR1 sample was reinjected with targeted MS^2^ and MS^3^ data acquisition for the *m/z* 397.1348 analyte, and the results are shown in Figure 3 and tabulated in Table S4. As indicated in Table 1, the putative molecular formula is C_16_H_20_N_4_O_8,_ differing from riboflavin (C_17_H_20_N_4_O_6_) by two additional oxygen atoms and one fewer carbon atom.

**Table 1.** Ribityl derivative analytes in decreasing order of intensity with >5 fold higher levels in the MR1 compared to CD1 and linked together in the molecular network cluster with putative molecular formulas, modifications and metabolism transformations based upon the “Generated Expected Compounds” feature of CD and the molecular structure of riboflavin. Analyte with a mass of 231.1474 Da was excluded from the table because the fragmentation spectra and mass are not consistent with the compound class.(a) The *m/z* 419.1180 was observed and attributed to the sodiated analyte. (b) Analytes found in Harriff et al.(23) (c) Analytes found within the cell media. (d) Confirmed with spiked-in ^13^C_4_,^15^N_2_-riboflavin and comparison of fragmentation data with synthetic standard. (e) The *m/z* 414.1160 was observed and attributed to the sodiated analyte. (f) Molecular formula consistent with oxidation or dealkylation with desaturation of riboflavin. (g) Molecular formula consistent with desaturation, nitro reduction and glycine conjugation of riboflavin. (h) Relative abundance of each analyte is approximated as the area of the chromatographic peak as expressed as n x 10^9^.

| RT (min) | Molecular Weight (Da) | Putative Molecular Formula | Putative Modification | Max. Area (n x 10<sup>9</sup>)<sup>h</sup> | $\Delta$ Mass (ppm) |
| --- | --- | --- | --- | --- | --- |
| 20.5 | 396.1285<sup>a</sup> | C<sub>16</sub>H<sub>20</sub>N<sub>4</sub>O<sub>8</sub> | +O<sub>2</sub>, -C | 7.1 | 2.3 |
| **23.3** | **376.1385<sup>b,c</sup>** | **C<sub>17</sub>H<sub>20</sub>N<sub>4</sub>O<sub>6</sub>** | **Riboflavin<sup>d</sup>** | **1.5** | **2.1** |
| 25.8 | 420.1110<sup>b</sup> | C<sub>18</sub>H<sub>20</sub>N<sub>4</sub>O<sub>6</sub>S | +CS | 0.60 | 1.5 |
| 26.1, 21.3 | 392.1337<sup>b,c,e,f</sup> | C<sub>17</sub>H<sub>20</sub>N<sub>4</sub>O<sub>7</sub> | +O | 0.27, 0.11 | 1.4 |
| 26.0 | 387.1549 | C<sub>18</sub>H<sub>21</sub>N<sub>5</sub>O<sub>5</sub> | +CNH, -O | 0.19 | 1.5 |
| 26.2 | 431.1810 | C<sub>20</sub>H<sub>25</sub>N<sub>5</sub>O<sub>6</sub> | +C<sub>3</sub>H<sub>5</sub>N | 0.19 | 1.2 |
| 20.9 | 407.1085 | C<sub>16</sub>H<sub>17</sub>N<sub>5</sub>O<sub>8</sub> | +NO<sub>2</sub>, -CH<sub>3</sub> | 0.16 | 2.1 |
| 35.1 | 443.2176<sup>c</sup> | C<sub>22</sub>H<sub>29</sub>N<sub>5</sub>O<sub>5</sub> | +C<sub>5</sub>H<sub>9</sub>N, -O | 0.16 | 1.5 |
| 27.0 | 401.1704<sup>g</sup> | C<sub>19</sub>H<sub>23</sub>N<sub>5</sub>O<sub>5</sub> | +C<sub>2</sub>H<sub>3</sub>N, -O | 0.15 | 1.2 |
| 24.8 | 417.1654 | C<sub>19</sub>H<sub>23</sub>N<sub>5</sub>O<sub>6</sub> | +C<sub>2</sub>H<sub>3</sub>N | 0.12 | 1.5 |
| 21.6, 23.5 | 378.1181<sup>b</sup> | C<sub>16</sub>H<sub>18</sub>N<sub>4</sub>O<sub>7</sub> | +O, -CH<sub>2</sub> | 0.05, 0.04 | 1.6 |
| 26.8 | 380.1339 | C<sub>16</sub>H<sub>20</sub>N<sub>4</sub>O<sub>7</sub> | +O, -C | 0.03 | 1.8 |

**Figure 3.**
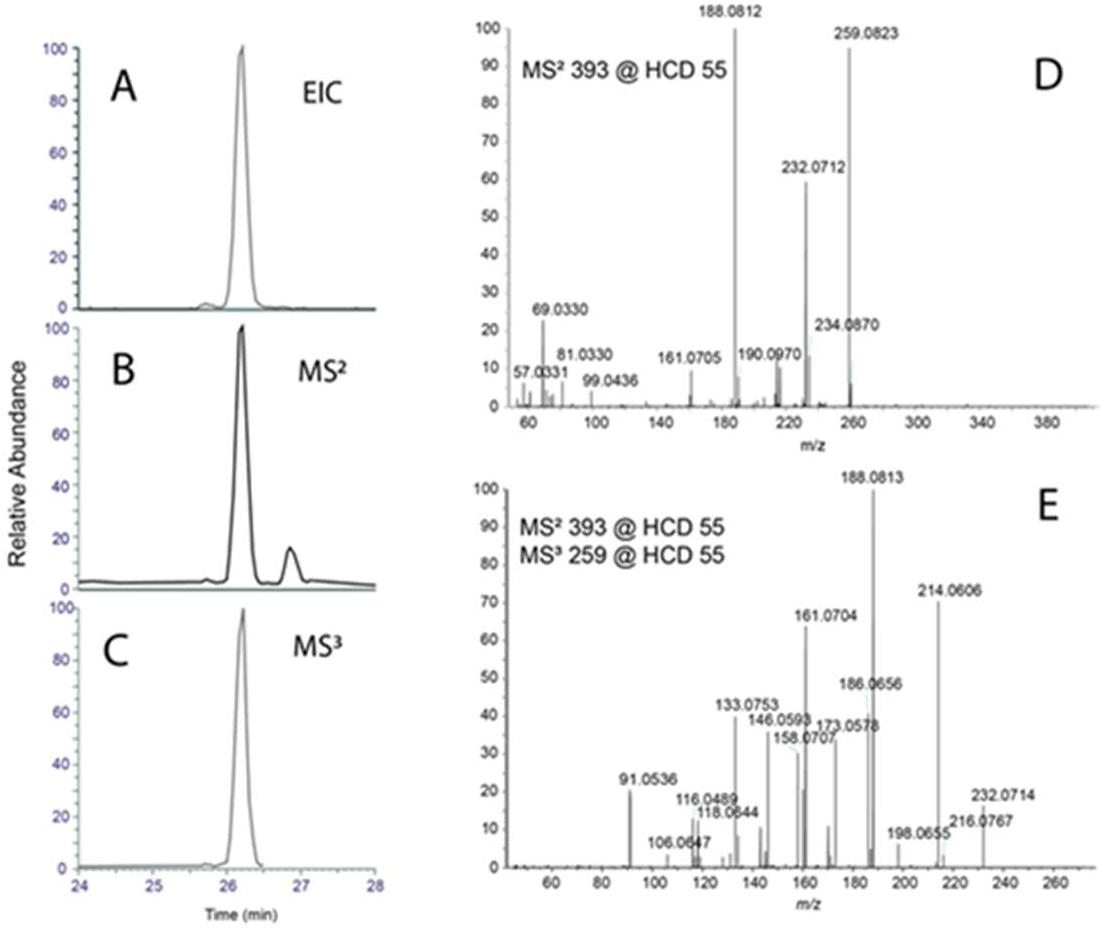
Chromatograms and corresponding spectra of the unknown *m/z* 397.1348 analyte. Panel A) Extracted precursor ion chromatogram from the full scan. Panel B) Base peak chromatogram for the *m/z* 397 MS^2^ spectra. Panel C) Corresponding MS^2^ spectrum Panel E) Corresponding MS^3^ spectrum.

Studies reporting ribityl-containing analytes in microbial samples were conducted using negative ionization mode,^15, 24, 25^ in contrast to the positive ion mode data presented here. The analytes in Table 1 with molecular masses of 378.1181 and 392.1337, along with riboflavin (376.1385 Da), were noted by Harriff et al^24^ as being assigned as 8- demethyl-8-hydroxy-riboflavin and 6-hydroxy-riboflavin, respectively, based on *m/z* values, isotope distributions, unsaturation counts, published literature, and similarities in fragment ion spectra/structures with known clusters within a molecular network. A manual interrogation of the data for all previously reported sMR1 ligands^15, 24, 25^ also revealed a precursor mass within 5 ppm of acetyl RL-6-Me-7-OH (370.1124 Da, *m/z* 371.1197) exclusively in the sMR1 samples. However, the fragmentation spectrum (Figure S4) exhibited a dominant product ion consistent with a ribityl moiety, which is absent in the chemical structure of acetyl RL-6-Me-7-OH. This suggests an additional newly identified putative riboflavin-related sMR1 ligand with an extracted precursor ion chromatographic peak area of 2.4e8, corresponding to a signal intensity that is similar to, but lower than, the lowest abundant ion of the novel ribityl-related compounds observed here and summarized in Table 1.

Serum-free cell media used in the sMR1 and sCD1d experiments was processed and analyzed using our DDA-CNL/MS^3^ analysis to determine if the analytes listed in Table 1 were present in the cell media. As expected, riboflavin was present with intense ion signals consistent with those observed in the sMR1 and sCD1d samples. Trace levels of signals consistent with the 421.1360 Da, 444.2239 Da, and 393.1405 Da analytes were also observed; however, the signal intensities were 0.015%, 0.025%, and 0.15% of that of riboflavin, with the 393.1405 Da analyte present at similar levels in a blank injection.

### Identification of Adenine-Containing Compounds

We then used our CD data analysis workflow to investigate the dataset for additional MR1-binding compounds using the CD “Neutral Loss” and the “Compound Class” nodes with the MS^2^ neutral losses and/or product ions, respectively, found in Table S1. We identified 462 analytes (features), in contrast to the ribityl-loss-only results, where 73 analytes were observed. Further filtering for a signal intensity sMR1/sCD1d ratio greater than 5, along with the neutral loss/compound class criteria, resulted in 85 analytes.

### Molecular Networking for Adenine-Containing Compounds

A CD molecular networking analysis (Figure 4) was employed to characterize the 85 analytes by clustering them based on their similar chemical structures. The ribityl cluster was again present (Figure 4, Panel A). A second cluster was observed, featuring multiple nodes defined by the mzCloud similarity matching feature of CD as being structurally related to adenosine and/or adenine (Figure 4, Panel B). This includes the putative identification of *m/z* 348.0708 as either 3’-adenosine monophosphate or adenosine 5’- monophosphate (347.0635 Da, AMP), based on the NIST 2020 MSMS HR database precursor mass and matching fragmentation spectra (Figure 5). A second discrete analyte, consistent with the general structure of AMP, was observed at similar levels in the sMR1s and sCD1d samples. The most abundant analytes in the cluster (Figure 4, Panel B) are listed in Table 2, while the complete list can be found in Table S3. The highest signal intensities for the analytes in Table 2 are plotted in Figure S5, with the analyte putatively assigned to AMP showing the most intensity.

**Table 2.**
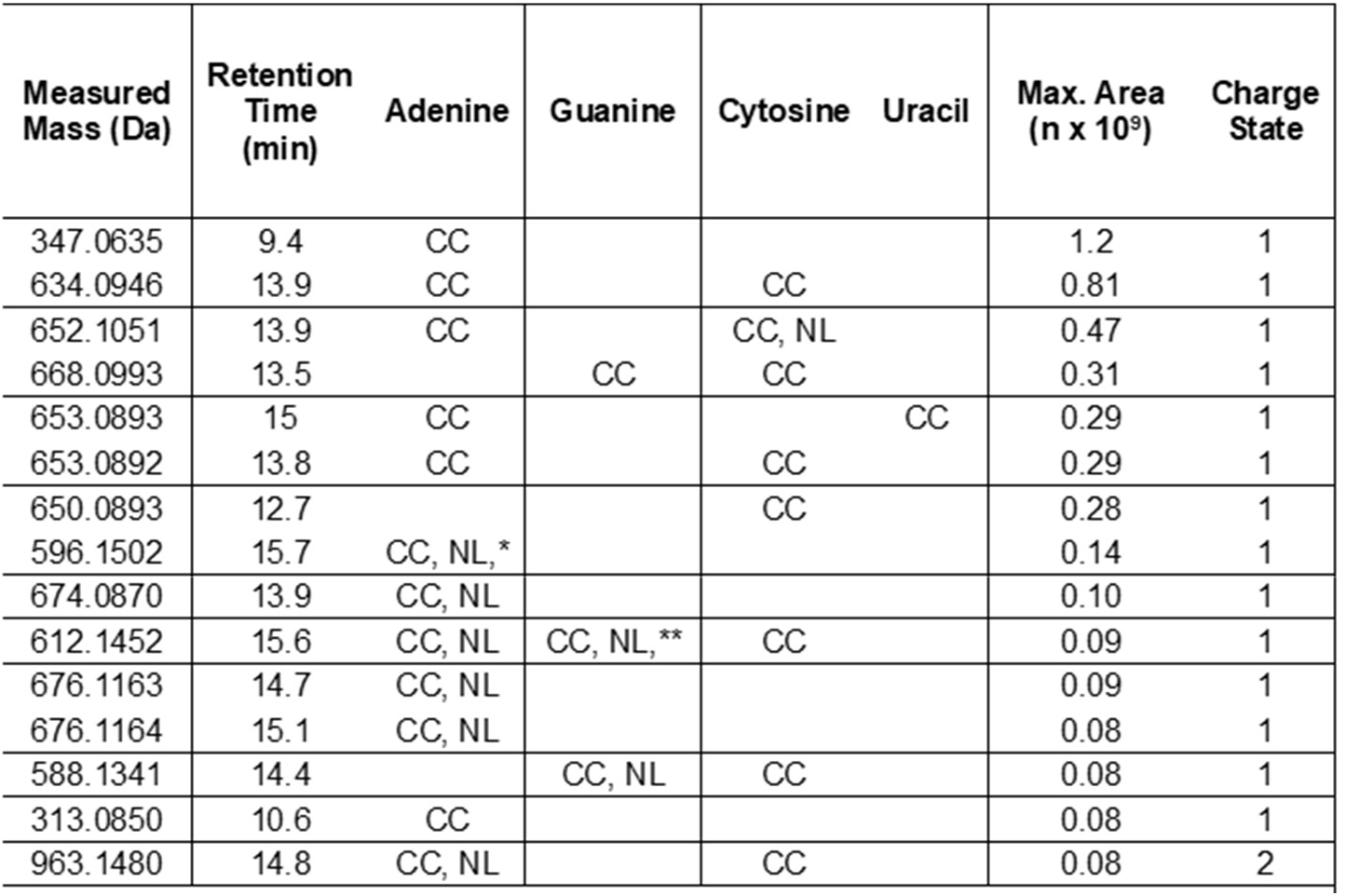
Compounds identified in the adenine-based cluster with CC indicating observation via the Compound Class node as a protonated nucleobase and NL indicating observation of the neutral loss of one of the nucleobases. The primary charge state observed is indicated as either “1” or “2” indicating detection as [M+1H]*^1^ and [M+2H]*^2^, respectively. The * and ** symbols indicate observation of the neutral loss of adenosine and guanosine, respectively.

**Figure 4.**
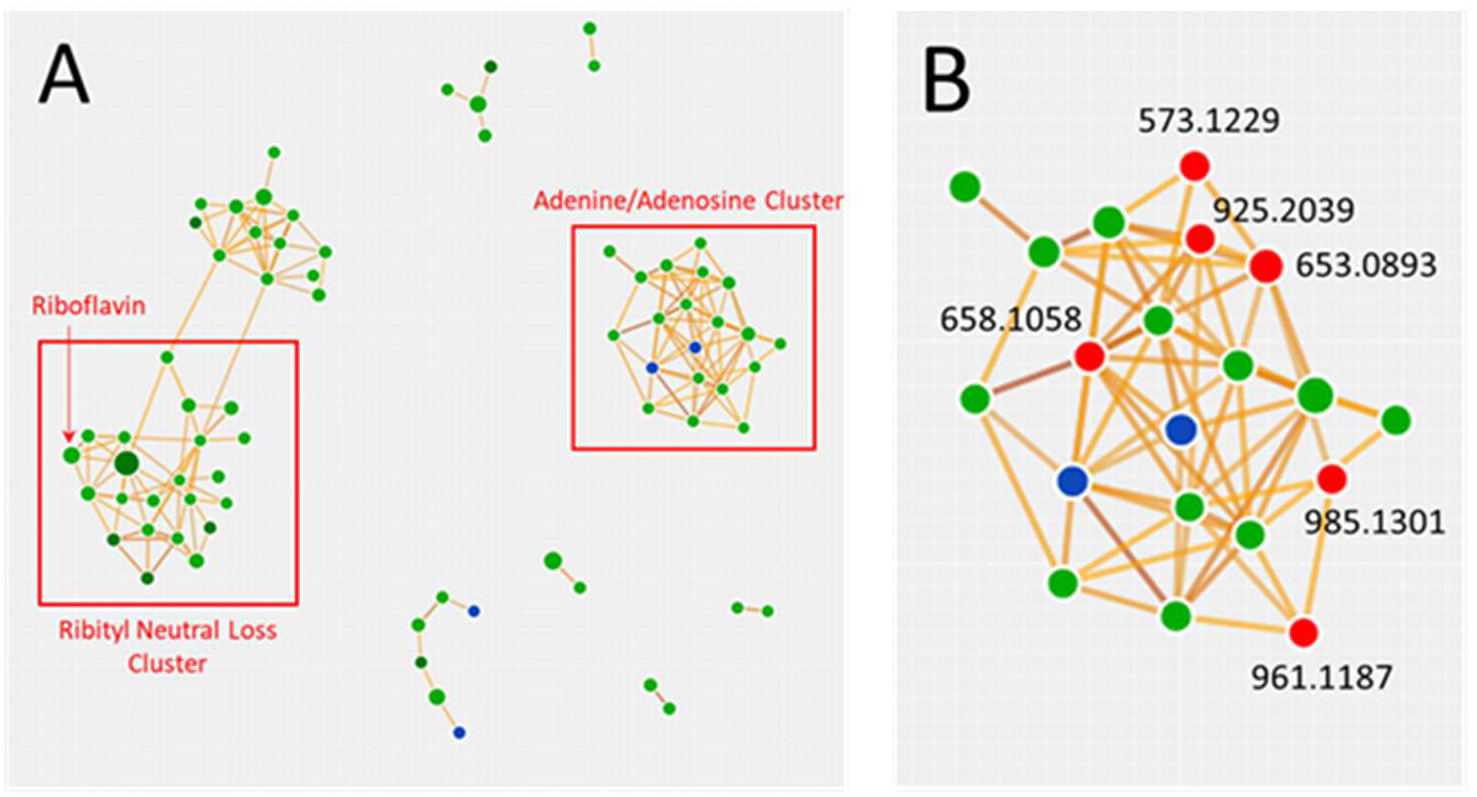
Molecular networking map without orphans shown and thresholds of score = 40, coverage = 60 and matched fragments = 3. Panel A) Full molecular network for those analytes with an sMR1/sCD1d ratio > 5 and a neutral loss/compound class found Panel B) Expanded region of adenine/adenosine cluster Red nodes correspond to those analytes which were identified as related to adenosine.

**Figure 5.**
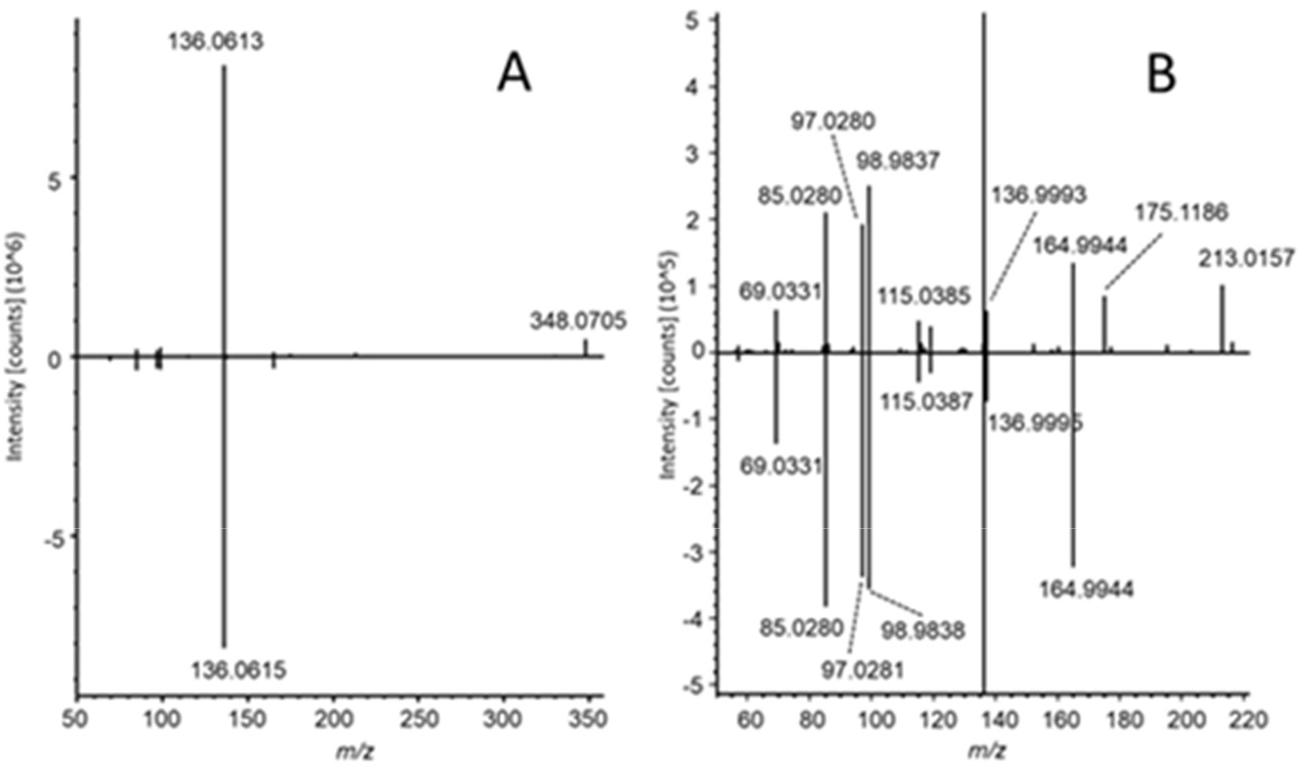
Panel A is the mirror plot of the fragmentation spectrum (stepped HCD of 25 and 50%) of the m/z 348 0708 feature at 9.4 minutes (top) and the spectrum (HCD 45%) for adenosine 3’- monophosphate entry in the NIST 2020 MSMS HR database (bottom). Panel B is the expanded range of Panel A.

None of the analytes listed in Table 2 were detected in the analyzed cell media samples. However, a loss of AMP due to the sample purification through solid-phase extraction, which is necessary for MS analysis, cannot be excluded.

### Structural Characterization of Adenine Phosphate Compound

The identification of AMP and other adenosine- related compounds suggests that adenosine monophosphate groups are a common structural subunit of the significantly larger mass nodes detected (*m/z* 529 - 1293). All four RNA bases are observed in various fragmentation spectra following the frequency order of adenine>cytosine>guanine>uracil, indicating the presence of multiple RNA nucleotides as substructures of the analytes in the cluster. Of the 33 analytes identified, 30 contained adenine as a neutral loss and/or a product ion, suggesting that AMP is the dominant substructure. The two analytes with the highest ion abundances, 634.0946 Da and 652.1050 Da, display MS^2^ spectra indicating the presence of adenine and cytosine, with masses consistent with AMP and cytosine monophosphate (CMP) covalently bound to each other, e. g., AMP+CMP-H_2_O and AMP+CMP, respectively. The structural information gained from the MS^2^ and MS^3^ spectra cannot be used to identify structures. However, the masses of many of the analytes and their fragmentation spectra are consistent with the covalent linking of combinations of various nucleotides. An example of this is the *m/z* 677.1229 (676.1163 Da) analyte, which has a mass consistent with the covalent binding of two AMP units and a clean, relatively simple fragmentation spectrum (Figure 6), dominated by the product ions of protonated adenine and AMP and the neutral loss of adenine.

**Figure 6.**
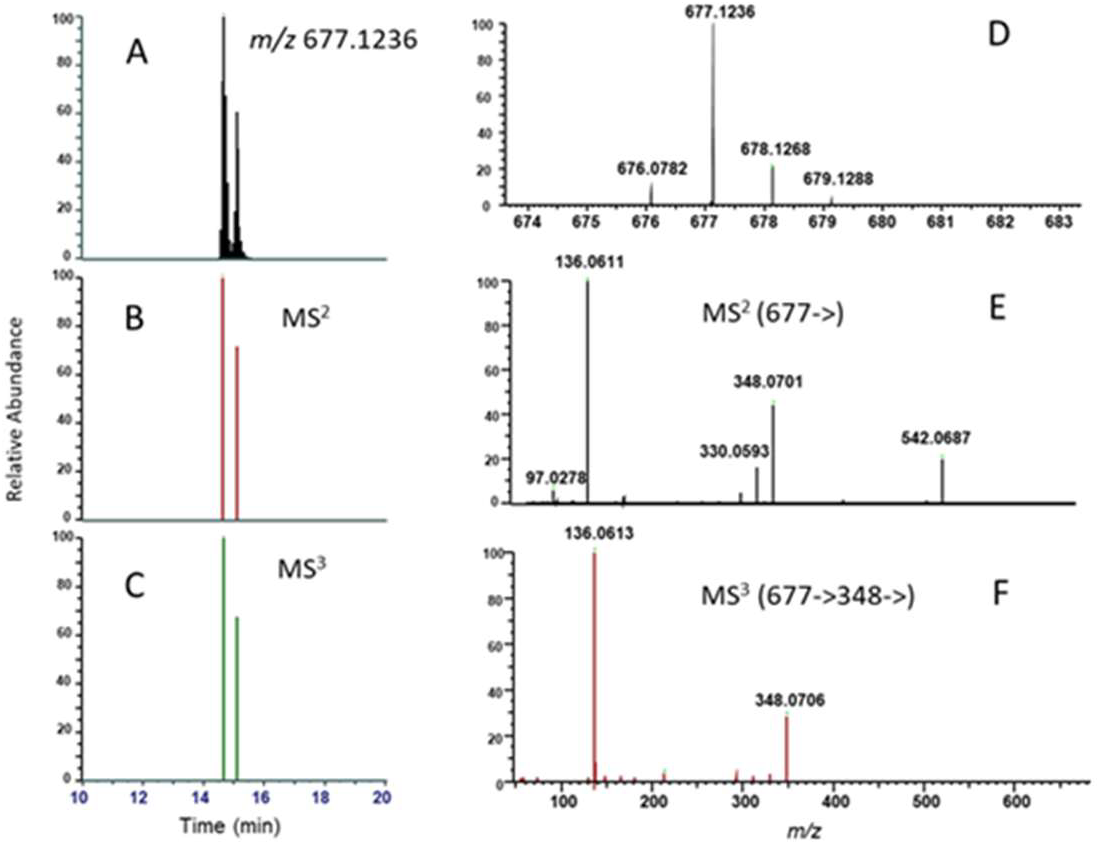
Representative chromatogram and spectra of analyte identified by appearance of protonated adenine and neutral loss of adenine. Panel A) Extracted *m/z* 677.1236 signal from the full scan. Panel B) Data dependent MS^2^ event for the 677.1236 ion. Panel C) MS^3^ signal triggered by the detection of the protonated adenine ion in the MS^2^ spectrum. Panel D) Full scan signal for the 677.1236 analyte corresponding to Panel A. Panel E) The MS^2^ spectrum from Panel B Panel F)MS^3^ triggered spectrum from Panel C.

One possible structure for this analyte is the protonated di-adenine RNA dinucleotide (ApAp), which was purchased as a synthetic standard for comparison to the unknown. A second possibility is the P^1^P^2^-diadenosine-5’- pyrophosphate (AppA) compound, which is structurally similar to nicotinamide adenine dinucleotide (NAD). This compound was synthesized using a method previously reported in the literature^21^ and described in the Supporting Information. Chemical structures for AppA and ApAp are shown in Figure S6. These two synthetic standards were injected using the same LC-MS method employed to collect the sMR1 and sCD1d data, and their chromatography and fragmentation spectra are compared to those of the *m/z* 677.1229 analyte (Figure 7). This comparison revealed a dominant protonated adenine ion in the MS^2^ spectra of all three, but apparent overall differences in the MS^2^ spectra and retention times of the synthetic standards and the analyte, indicating the analyte is neither of the two synthetic standards.

**Figure 7.**
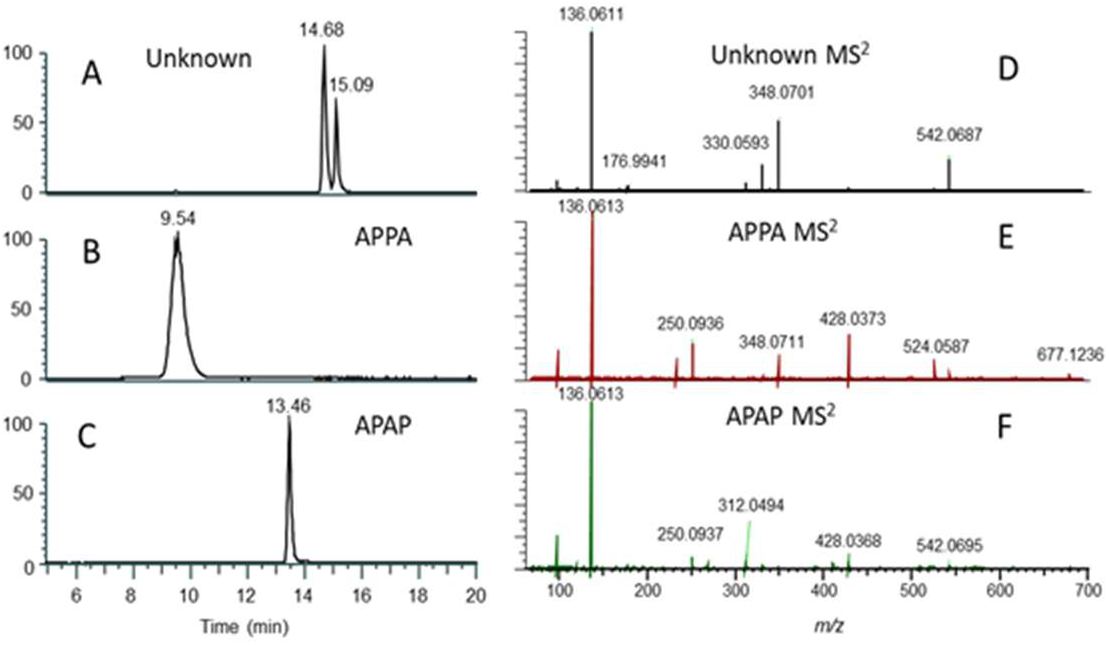
Comparison between the unknown analyte found in the sMR1 and commercial ApAp and synthetic AppA standard.

### Identified sMR1-ligand Compound Classes of Mammalian Origin

Classes of compounds share structural moieties and functional groups that, upon MS^n^ fragmentation, often result in diagnostic product ions and/or neutral losses. These fragmentation patterns can be used to screen samples for known and unknown analytes of specific classes of compounds. We have previously employed this concept to aid in the identification of modified nucleosides in a study of ligands bound to MR1.^12^ We reanalyzed these data-dependent acquisition datasets to probe for additional classes of compounds within the sMR1 ligandome using Compound Discoverer, a commercial software package designed for metabolomic analysis, with its fragmentation filtering nodes (Neutral Loss and Compound Class), molecular networking, and searching of mzCloud and NIST 2020 HR spectral databases. This structure-oriented LC-MS discovery data analysis can also be performed using freeware such as MZMINE^26^ (with the added functionality of the DFBuilder algorithm^27^), MS-FINDER,^28^ XCMS,^29^ MS-DIAL,^30^ GNPS,^19^ and Spec2Vec.^31^

Our CD analysis identified two compound classes with significantly higher levels in sMR1 samples compared to sCD1d samples, another nonclassical antigen-presenting molecule. One class consists of ribityl-containing compounds, some or all of which appear to be structurally similar to riboflavin. The mass difference between the most abundant (*m/z* 396.1285, 66%) of these analytes and riboflavin (*m/z* 376.1385, 14%) is 19.9900 Da, which is consistent with a molecular formula difference of one less carbon atom and two additional oxygen atoms. It is not attributed to Phase I and/or Phase 2 metabolism (considering up to two Phase 1 and one Phase 2 transformations) or found in BioCyc or Metabolika. The identity of this analyte and the lower abundance ribityl-containing analytes remains an open question. The MS^2^ and MS^3^ spectra of the *m/z* 396.1285 analyte are reported here in Figure 3 to aid future efforts in identifying its structure. A recent review^17^ of MR1 ligands listed three classes of vitamin B (2, 6, and 9) compounds as antigens, including ribityl-containing riboflavin and related vitamin B2 compounds which are discussed above. Of the listed vitamin B9 and B6 related antigens, the vitamin B9 folic acid and the recently found^18^ vitamin B6 pyridoxol are both observed in our dataset and identified through molecular mass and fragmentation spectral matching to mzCloud database entrees. The extracted precursor ion chromatographic peak areas of pyridoxol, folic acid, and riboflavin are approximately 6, 10, and 8 times higher in the sMR1 as compared to the sCD1 samples, respectively. The maximum peak areas observed for pyridoxol, folic acid, and riboflavin are 0.88e9, 0.41e9, and 1.50e9, respectively, and the mzCloud ‘Best Match” values are 84.9, 85.5, and 80.2, respectively (Figure S7).

The other class of compounds elevated in the sMR1 samples consists of combinations of nucleobases, ribose groups, and phosphate groups, including adenosine monophosphate (AMP). The two most abundant analytes in this category have masses and fragmentation spectra that are consistent with covalently bound adenosine and cytosine monophosphate groups. A series of less abundant analytes is comprised of nucleobases, ribose, and phosphate groups covalently linked to one another, with adenine being the dominant nucleobase. To understand the nature of this compound class, the chromatographic and fragmentation spectra of the *m/z* 677.1229 analyte, which has a putative molecular formula consistent with two covalently bound adenosine monophosphate molecules, were compared to synthetic standards of AppA and ApAp, revealing apparent differences between them and the unknown analyte. A third possible structure comprises two adenine monophosphate groups bound through their ribose groups (Figure S6), similar to the polyadenosine 5’-diphosphoribose structures produced in the PARylation processes^32, 33^ The nature of this class of compounds requires further investigation. The ability to link structurally similar compounds to known chemical structures and to identify structural subunits is a powerful tool for discovering novel classes of compounds. However, confidently assigning a molecular identity to the newly discovered analytes remains challenging, as demonstrated by our inability to match one of the analytes to two plausible synthetic standards. This example highlights the caution needed in assigning chemical structures based on MS^n^ data.

The ability to predict either molecular structures from fragmentation spectra or fragmentation spectra from molecular structures is a challenging task with many ongoing efforts and varying levels of success.^34, 35^ The MS^2^ and MS^3^ spectra for the 392.1337 Da and 676.1163 Da analytes reported here can be used to putatively identify these compounds with the ever-expanding set of analytical tools. Despite the challenges of analyte structural characterization, the approaches used here are powerful tools for identifying novel classes of compounds, investigating relevant biological transformations, and characterizing novel processes and pathways. A recent approach utilizing known biotransformation rules, sourced from databases such as RetroRules,^36^ KEGG,^37^ or Metacyc,^38^ could be incorporated into the workflow reported here, with riboflavin and AMP serving as anchors to characterize the chemical structures of the observed unknowns.^39^

## CONCLUSION

This is the first example of applying our HRMS-DDA-MS^3^ data acquisition methodology and data analysis workflow to identify unknown molecules produced by mammalian cells that may have important physiological roles. The characterization of the MR1 ligandome extends the DNA adductomics workflow, traditionally used to characterize DNA damage, to explore novel classes of compounds and uncover new pathways with potential immunological relevance. These results demonstrate the power of combining compound-class fragmentation with molecular networking and the searching of mass spectral databases.

## ASSOCIATED CONTENT

### Data Availability Statement

#### Supporting Information

Scheme S1 - Synthesis of AppA

Synthesis of AppA - detailed description of synthesis and characterization by NMR

Figure S1 - ^1^H-NMR of synthetic AppA

Figure S2 - ^31^P-NMR of synthetic AppA

Table S1 - Neutral losses and product ions associated with nucleobases

Table S2 - List of precursor ions for targeted DDA-CNL/MS3 detection at high HCD energy

Scheme S2 - Compound Discoverer graphical workflow

Compound Discoverer workflow parameters

Table S3 - Peak intensities for the nucleobase phosphate analytes

Table S4 - Product ions and relative abundances for MS^2^ fragmentation of *m/z* 397.1358 analyte

Figure S3 - Plot of peak intensities of ribityl analytes

Figure S4 - MS detection of 370.1124 Da analyte

Figure S5 - Plot of peak areas of most abundant nucleobase phosphate analytes

Figure S6 - Possible structures for the 676.1163 Da analyte

Figure S7 - Extracted ion chromatograms and MS^2^ mirror plots of pyridoxal, folic acid and riboflavin

## AUTHOR INFORMATION

## Authors

**Chiara Lecchi** - *Masonic Cancer Center, University of Minnesota; Minneapolis, MN 55455, United States*;

**Alessandro Vacchini** - *Experimental Immunology, Department of Biomedicine, University Hospital and University of Basel, Basel 4031, Switzerland;*

**Stefano Sainas** - *Department of Drug Science and Technology. University of Torino, Via Pietro Giuria 9, Torino 10125, Italy*;

**Marco L. Lolli** - *Department of Drug Science and Technology. University of Torino, Via Pietro Giuria 9, Torino 10125, Italy*;

**Matthew Luedtke** *- Masonic Cancer Center, University of Minnesota; Minneapolis, MN 55455, United States*;

**Lucia Mori** - *Experimental Immunology, Department of Biomedicine, University Hospital and University of Basel, Basel 4031, Switzerland;*

## Author Contribution

GDI, LM, and AV conceived the study with subsequent conceptual input from SB and PWV. PWV, CL, AV, SS, MLL and ML performed the investigation. GDL and SB acquired funding. PWV and SB wrote the original draft. All authors reviewed, edited, and approved the manuscript.

## ACKNOWLEDGEMENTS

This work was supported by grants from the Swiss National Foundation 310030-173240 (G.D.L.), Swiss Cancer League KFS-4707-02-2019 (G.D.L.), Basel Cancer League KLbB-4779-02-2019 (G.D.L.), D-BSSE ETH Zürich PMB-02-17 (G.D.L.), and University Hospital of Basel (G.D.L.); by University of Turin, Ricerca Locale (SAIS_RILO_24_01), by Matterhorn Biosciences AG (G.D.L. and S.B.); and the NIH National Cancer Institute R50-CA211256 (P.W.V). Mass spectrometry was performed at the University of Minnesota, Masonic Cancer Center, Analytical Biochemistry Shared Resource partly funded by NIH National Cancer Institute grant P30CA077598.

## REFERENCES

1. Purcell, A. W., Ramarathinam, S. H., and Ternette, N.: Mass spectrometry-based identification of MHC-bound peptides for immunopeptidomics. Nat Protoc 14, 1687–1707 (2019)

2. Chong, C., Coukos, G., and Bassani-Sternberg, M.: Identification of tumor antigens with immunopeptidomics. Nat Biotechnol 40, 175–188 (2022)

3. Bumpus, S. B., Evans, B. S., Thomas, P. M., Ntai, I., and Kelleher, N. L.: A proteomics approach to discovering natural products and their biosynthetic pathways. Nat Biotechnol 27, 951–956 (2009)

4. Lai, Y., Koelmel, J. P., Walker, D. I., Price, E. J., Papazian, S., Manz, K. E., et al.: High-Resolution Mass Spectrometry for Human Exposomics: Expanding Chemical Space Coverage. Environ Sci Technol 58, 12784–12822 (2024)

5. Beger, R. D., Goodacre, R., Jones, C. M., Lippa, K. A., Mayboroda, O. A., O’Neill, D., et al.: Analysis types and quantification methods applied in UHPLC-MS metabolomics research: a tutorial. Metabolomics 20, 95 (2024)

6. Tretyakova, N., Villalta, P. W., and Kotapati, S.: Mass spectrometry of structurally modified DNA. Chem Rev 113, 2395–2436 (2013)

7. Carrasco, M., and Mirzaei, H. (2016) Modern Proteomics - Sample Preparation, Analysis and Practical Applications, In Advances in Experimental Medicine and Biology, pp 1 online resource (IX, 533 pages 180 illustrations, 130 illustrations in color, Springer International Publishing : Imprint: Springer,, Cham.

8. Jiang, Y., Stornetta, A., Villalta, P. W., Wilson, M. R., Boudreau, P. D., Zha, L., et al.: Reactivity of an Unusual Amidase May Explain Colibactin’s DNA Cross-Linking Activity. J Am Chem Soc 141, 11489–11496 (2019)

9. Stornetta, A., Villalta, P. W., Hecht, S. S., Sturla, S. J., and Balbo, S.: Screening for DNA Alkylation Mono and Cross-Linked Adducts with a Comprehensive LC-MS(3) Adductomic Approach. Anal Chem 87, 11706–11713 (2015)

10. Ragi, N., Walmsley, S. J., Jacobs, F. C., Rosenquist, T. A., Sidorenko, V. S., Yao, L., et al.: Screening DNA Damage in the Rat Kidney and Liver by Untargeted DNA Adductomics. Chem Res Toxicol 37, 340–360 (2024)

11. Walmsley, S. J., Guo, J., Tarifa, A., DeCaprio, A. P., Cooke, M. S., Turesky, R. J., and Villalta, P. W.: Mass Spectral Library for DNA Adductomics. Chemical Research in Toxicology 37, 302–310 (2024)

12. Vacchini, A., Chancellor, A., Yang, Q., Colombo, R., Spagnuolo, J., Berloffa, G., et al.: Nucleobase adducts bind MR1 and stimulate MR1-restricted T cells. Sci Immunol 9, eadn0126 (2024)

13. Kjer-Nielsen, L., Patel, O., Corbett, A. J., Le Nours, J., Meehan, B., Liu, L., et al.: MR1 presents microbial vitamin B metabolites to MAIT cells. Nature 491, 717–723 (2012)

14. Corbett, A. J., Eckle, S. B., Birkinshaw, R. W., Liu, L., Patel, O., Mahony, J., et al.: T-cell activation by transitory neo-antigens derived from distinct microbial pathways. Nature 509, 361–365 (2014)

15. McWilliam, H. E. G., and Villadangos, J. A.: MR1 antigen presentation to MAIT cells and other MR1-restricted T cells. Nat Rev Immunol 24, 178–192 (2024)

16. Awad, W., Ler, G. J. M., Xu, W., Keller, A. N., Mak, J. Y. W., Lim, X. Y., et al.: The molecular basis underpinning the potency and specificity of MAIT cell antigens. Nat Immunol 21, 400–411 (2020)

17. Awad, W., Abdelaal, M. R., Letoga, V., McCluskey, J., and Rossjohn, J.: Molecular Insights Into MR1-Mediated T Cell Immunity: Lessons Learned and Unanswered Questions. Immunol Rev 331, e70033 (2025)

18. McInerney, M. P., Awad, W., Souter, M. N. T., Kang, Y., Wang, C. J. H., Chan Yew Poa, K., et al.: MR1 presents vitamin B6-related compounds for recognition by MR1-reactive T cells. Proc Natl Acad Sci U S A 121, e2414792121 (2024)

19. Nothias, L. F., Petras, D., Schmid, R., Duhrkop, K., Rainer, J., Sarvepalli, A., et al.: Feature-based molecular networking in the GNPS analysis environment. Nat Methods 17, 905–908 (2020)

20. Morehouse, N. J., Clark, T. N., McMann, E. J., van Santen, J. A., Haeckl, F. P. J., Gray, C. A., and Linington, R. G.: Annotation of natural product compound families using molecular networking topology and structural similarity fingerprinting. Nat Commun 14, 308 (2023)

21. Ahmadibeni, Y., and Parang, K.: Solid-phase synthesis of symmetrical 5’,5’-dinucleoside mono-, di-, tri-, and tetraphosphodiesters. Org Lett 9, 4483–4486 (2007)

22. Lepore, M., Kalinichenko, A., Calogero, S., Kumar, P., Paleja, B., Schmaler, M., et al.: Functionally diverse human T cells recognize non-microbial antigens presented by MR1. Elife 6, (2017)

23. Du, Y., Li, Y. J., Hu, X. X., Deng, X., Qian, Z. T., Li, Z., et al.: Development and evaluation of a hydrophilic interaction liquid chromatography-MS/MS method to quantify 19 nucleobases and nucleosides in rat plasma. Biomed Chromatogr 31, (2017)

24. Harriff, M. J., McMurtrey, C., Froyd, C. A., Jin, H., Cansler, M., Null, M., et al.: MR1 displays the microbial metabolome driving selective MR1-restricted T cell receptor usage. Sci Immunol 3, (2018)

25. Krawic, J. R., Ladd, N. A., Cansler, M., McMurtrey, C., Devereaux, J., Worley, A., et al.: Multiple Isomers of Photolumazine V Bind MR1 and Differentially Activate MAIT Cells. J Immunol 212, 933–940 (2024)

26. Heuckeroth, S., Damiani, T., Smirnov, A., Mokshyna, O., Brungs, C., Korf, A., et al.: Reproducible mass spectrometry data processing and compound annotation in MZmine 3. Nat Protoc DOI: 10.1038/s41596-024-00996-y, (2024)

27. Murray, K. J., Carlson, E. S., Stornetta, A., Balskus, E. P., Villalta, P. W., and Balbo, S.: Extension of Diagnostic Fragmentation Filtering for Automated Discovery in DNA Adductomics. Anal Chem 93, 5754–5762 (2021)

28. Dai, W., Yin, P., Zeng, Z., Kong, H., Tong, H., Xu, Z., et al.: Nontargeted modification-specific metabolomics study based on liquid chromatography-high-resolution mass spectrometry. Anal Chem 86, 9146–9153 (2014)

29. Giera, M., Aisporna, A., Uritboonthai, W., Hoang, L., Derks, R. J. E., Joseph, K. M., et al.: XCMS-METLIN: data-driven metabolite, lipid, and chemical analysis. Mol Syst Biol 20, 1153–1155 (2024)

30. Tsugawa, H., Cajka, T., Kind, T., Ma, Y., Higgins, B., Ikeda, K., et al.: MS-DIAL: data-independent MS/MS deconvolution for comprehensive metabolome analysis. Nat Methods 12, 523–526 (2015)

31. Huber, F., Ridder, L., Verhoeven, S., Spaaks, J. H., Diblen, F., Rogers, S., and van der Hooft, J. J. J.: Spec2Vec: Improved mass spectral similarity scoring through learning of structural relationships. PLoS Comput Biol 17, e1008724 (2021)

32. Wei, H., and Yu, X.: Functions of PARylation in DNA Damage Repair Pathways. Genomics Proteomics Bioinformatics 14, 131–139 (2016)

33. Fabian, Z., Kakulidis, E. S., Hendriks, I. A., Kuhbacher, U., Larsen, N. B., Oliva-Santiago, M., et al.: PARP1-dependent DNA-protein crosslink repair. Nat Commun 15, 6641 (2024)

34. Xing, S., Hu, Y., Yin, Z., Liu, M., Tang, X., Fang, M., and Huan, T.: Retrieving and Utilizing Hypothetical Neutral Losses from Tandem Mass Spectra for Spectral Similarity Analysis and Unknown Metabolite Annotation. Anal Chem 92, 14476–14483 (2020)

35. Blazenovic, I., Kind, T., Ji, J., and Fiehn, O.: Software Tools and Approaches for Compound Identification of LC-MS/MS Data in Metabolomics. Metabolites 8, (2018)

36. Duigou, T., du Lac, M., Carbonell, P., and Faulon, J. L.: RetroRules: a database of reaction rules for engineering biology. Nucleic Acids Res 47, D1229–D1235 (2019)

37. Kanehisa, M., Furumichi, M., Sato, Y., Kawashima, M., and Ishiguro-Watanabe, M.: KEGG for taxonomy-based analysis of pathways and genomes. Nucleic Acids Res 51, D587–D592 (2023)

38. Balzerani, F., Blasco, T., Perez-Burillo, S., Valcarcel, L. V., Hassoun, S., and Planes, F. J.: Extending PROXIMAL to predict degradation pathways of phenolic compounds in the human gut microbiota. NPJ Syst Biol Appl 10, 56 (2024)

39. Martin, M. R., Bittremieux, W., and Hassoun, S.: Molecular Structure Discovery for Untargeted Metabolomics Using Biotransformation Rules and Global Molecular Networking. Anal Chem 97, 3213–3219 (2025)

