## Supplemental Material for "Expanding the MR1 ligandome using chemical class-specific fragmentation and molecular networking"

**SUPPORTING INFORMATION**

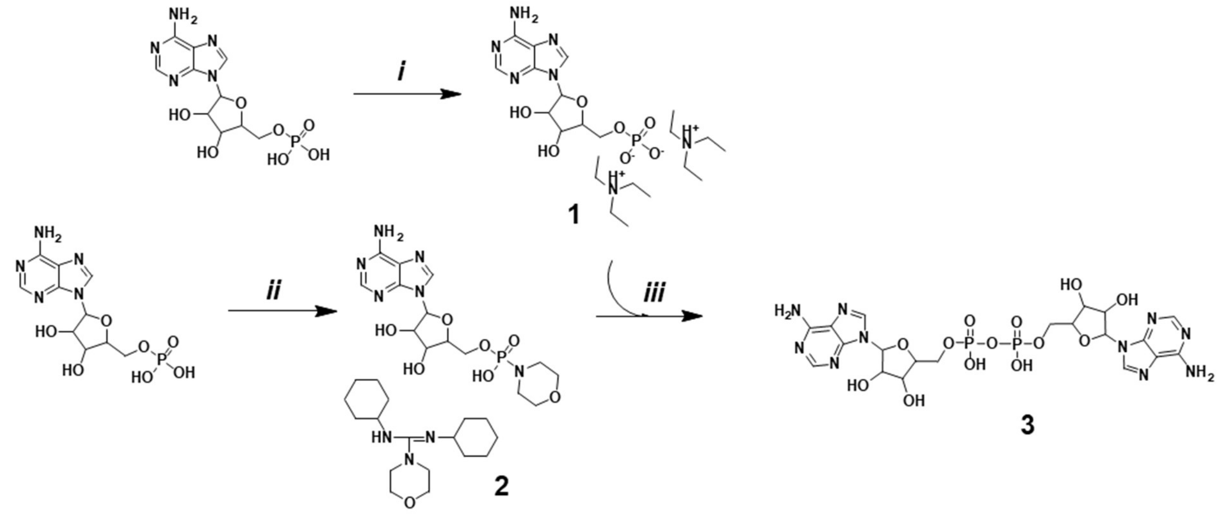


**Scheme S1.** *i*) dry $Et_3N$, dry MeOH; *ii*) Morpholine, DDC, 2-*But*OH; *iii*) dry pyridine.

### Synthesis of AppA

The synthesis and purification of AppA (**3**) were performed according to a previously published protocol(1, 2) and are described below.

*Bis (triethylammonium) salt 5'-Adenosinemonophosphate* (**1**). A solution of dry triethylamine (0.2 mmol, 30 µL) was added to a stirred suspension of 5'-adenosine-monophosphate free acid monohydrate (0.1 mmol, 150 mg) in 20 mL of dry methanol. The reaction mixture was stirred at room temperature for 30 minutes, after which a clear solution was obtained. The resulting solution was evaporated under reduced pressure. The resulting solid was dissolved in dry pyridine and evaporated under reduced pressure. This process was repeated four times, constantly in the presence of dry pyridine, to obtain the corresponding salt.

*Adenosine-5'-phosphoromorpholidate-4-morpholine-N, N'-dicyclohexylcarboxamidine Salt.* (**2**). A solution of dicyclohexylcarbodiimide (11.52 mmol, 2.38 g) in *t*-butyl alcohol (10 mL) was added dropwise to a refluxing solution of the adenosine-5' phosphate free acid (2.88 mmol, 1.0 g) in a mixture of water (20 mL), t-butyl alcohol (10 mL), and morpholine (11.52 mmol, 1.0 g). The addition was complete in 2 hours, and the reaction mixture was refluxed overnight until the reaction was complete. The reaction mixture was then cooled to room temperature, and the solid was filtered off. The filtrate was evaporated to remove the large amount of t-butyl alcohol, and the remaining aqueous phase was extracted with diethyl ether three times (10 mL each). The clear aqueous solution was evaporated to dryness under vacuum. The residue was taken up with 2 mL of methanol, and the addition of diethyl ether (10 mL) precipitated a sticky solid, which was triturated with fresh diethyl ether to afford the title compound as a white solid. Yield 24.5%.

*$P^1P^2$-diadenosine-5'-pyrophosphate* (AppA, **3**). Compound 2 (0.1 mmol, 100 mg) was dried through three evaporation steps of its solution in anhydrous pyridine. Separately, an equimolar concentration of compound 1 (0.1 mmol, 101 mg) was dissolved in 2 mL of anhydrous pyridine, and this compound was rendered anhydrous by three evaporation processes of its solution. The two pyridine solutions were mixed and dried through several evaporation steps until the solution became clear. Finally, the reaction was conducted by stirring at room temperature under a nitrogen atmosphere. The reaction was monitored by TLC (2-butano/water/acetic acid 5:3:2 v/v/v). After 96 hours, compound 2 disappeared, and the solvent was evaporated under reduced pressure. The crude material was dissolved in water, and the pH was adjusted to 8 with 1.0 N NaOH. The basic aqueous solution was extracted with diethyl ether (2 × 10 mL), and then the aqueous solution was evaporated under reduced pressure.

The final product, Ap2A, was purified by reversed-phase HPLC using a Waters Associates (Milford, MA) system equipped with a Shimadzu SPD-10A 0.2 mm Prep UV-vis detector set at 260 nm. Separation occurred on a Luna 5 μm C18(2) 100 Å 250 10 mm² column purchased from Phenomenex (Torrance, CA). A 38-minute method was developed using mobile phase A (10 mM ammonium formate, pH 4.0) and mobile phase B (50:50 MeOH and 10 mM ammonium formate, pH 4.0), with a constant flow rate of 4 mL/min. The mobile phase gradient started at 0% B and was maintained for 2 minutes, then increased to 40% B over 15 minutes, followed by a further increase to 90% B over 3 minutes, which was then held constant for 5 minutes. The instrument was then returned to 0% B and equilibrated for 10 minutes before the next injection. The product was collected as a white solid at a retention time of 14.5 minutes after solvent evaporation.

To remove residual formate salts, the product was first dissolved in 1 mL of 0.1 M NaOH. Then, a 20-minute method was developed using mobile phase A ($H_2O$) and mobile phase B (MeOH). The same column and flow rate were utilized as described in the previous paragraph. The mobile phase gradient started at 0% B, was maintained for 2 minutes, and was then increased to 100% B over 15 minutes. Finally, the gradient returned to 0% B over 2 minutes and was allowed to equilibrate for 10 minutes before the next injection. The product was collected at a retention time of 6.60 minutes and dried to yield a white solid.

The NMR spectra of Ap2A were recorded and interpreted on a Bruker 500 MHz spectrometer by dissolving the product in DMSO-d6. Chemical shifts are reported in parts per million (ppm). Residual solvent reference peaks for DMSO-d6 $^{1}H$ NMR are 2.50 ppm for DMSO-d6 and 3.33 ppm for residual $H_2O$. Peak splitting employs the following abbreviations: br s = broad singlet, s = singlet, d = doublet, and m = multiplet. The experimental 1D NMR spectral data aligns with literature values.(3) $^{1}H$ NMR (500 MHz, DMSO-$d_6$) $\delta$ 8.43 (s, 2H), 8.14 (s, 2H), 7.20 (s, 4H), 6.63 (br s, 2H), 5.91 (d, $J$ = 4.4 Hz, 2H), 5.26 (br s, 2H), 4.48 (m, 2H), 4.37 (m, 2H), 4.07 (m, 2H), 4.01 (m, 2H), 3.95 (m, 2H); $^{31}P$ NMR (202 MHz, DMSO-$d_6$) $\delta$-11.34; HRMS (Orbitrap Lumos): $[M + H]^+$ calc'd 677.1229; found 677.1217.

### REFERENCES


1. Ahmadibeni Y, Parang K. Solid-phase synthesis of symmetrical 5',5'-dinucleoside mono-, di-, tri-, and tetraphosphodiesters. Org Lett. 2007;9(22):4483-6.
2. Millo EZ, E.; Galatini, A.; Benatti, U.; Damonte, G. Simple Synthesis of P1P2-Diadenosine 5'-Pyrophosphate. Synthetic Communications. 2008;38(19):3260-9.
3. Yousef Ahmadibeni and Keykavous Parang. Solid-Phase Synthesis of Symmetrical 5',5'-Dinucleoside Mono-, Di-, Tri-, and Tetraphosphodiesters. *Org. Lett.* **2007**, *9*, 4483-4486.

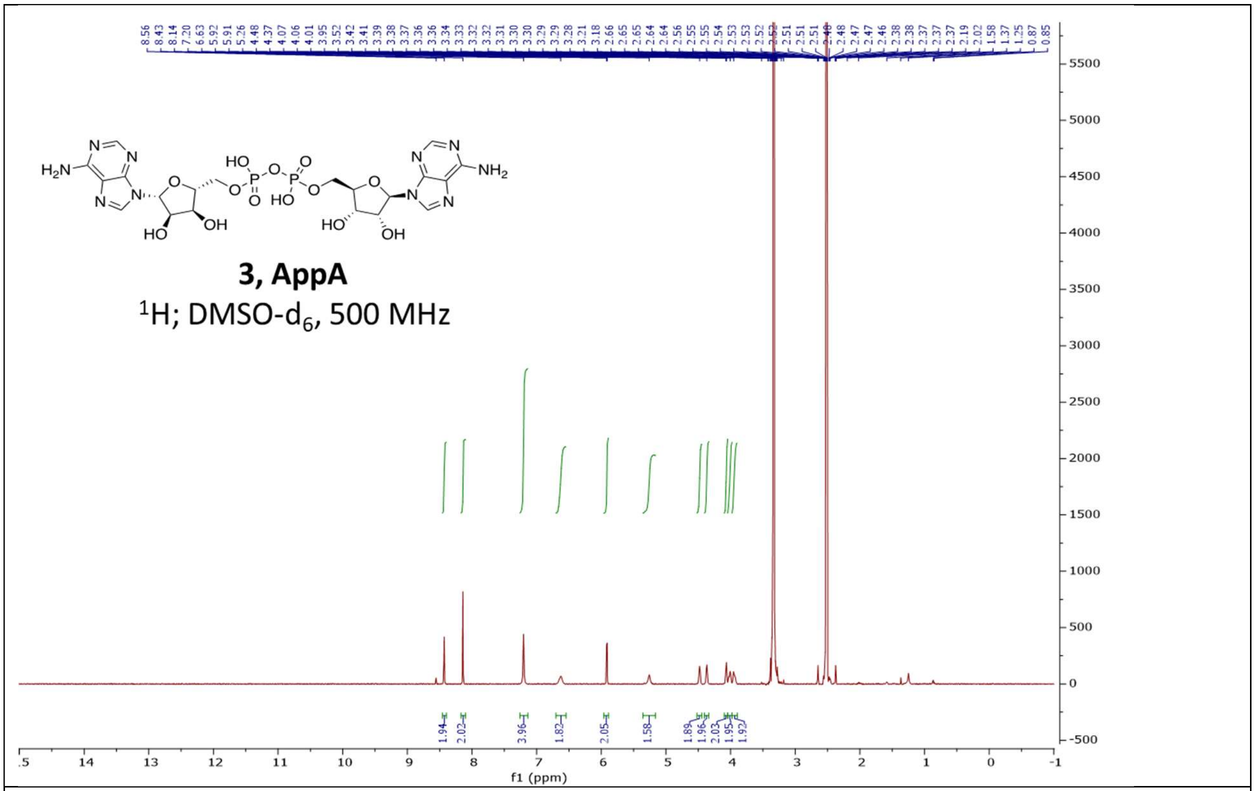


Figure S1. $^{1}$H-NMR of AppA synthetic standard

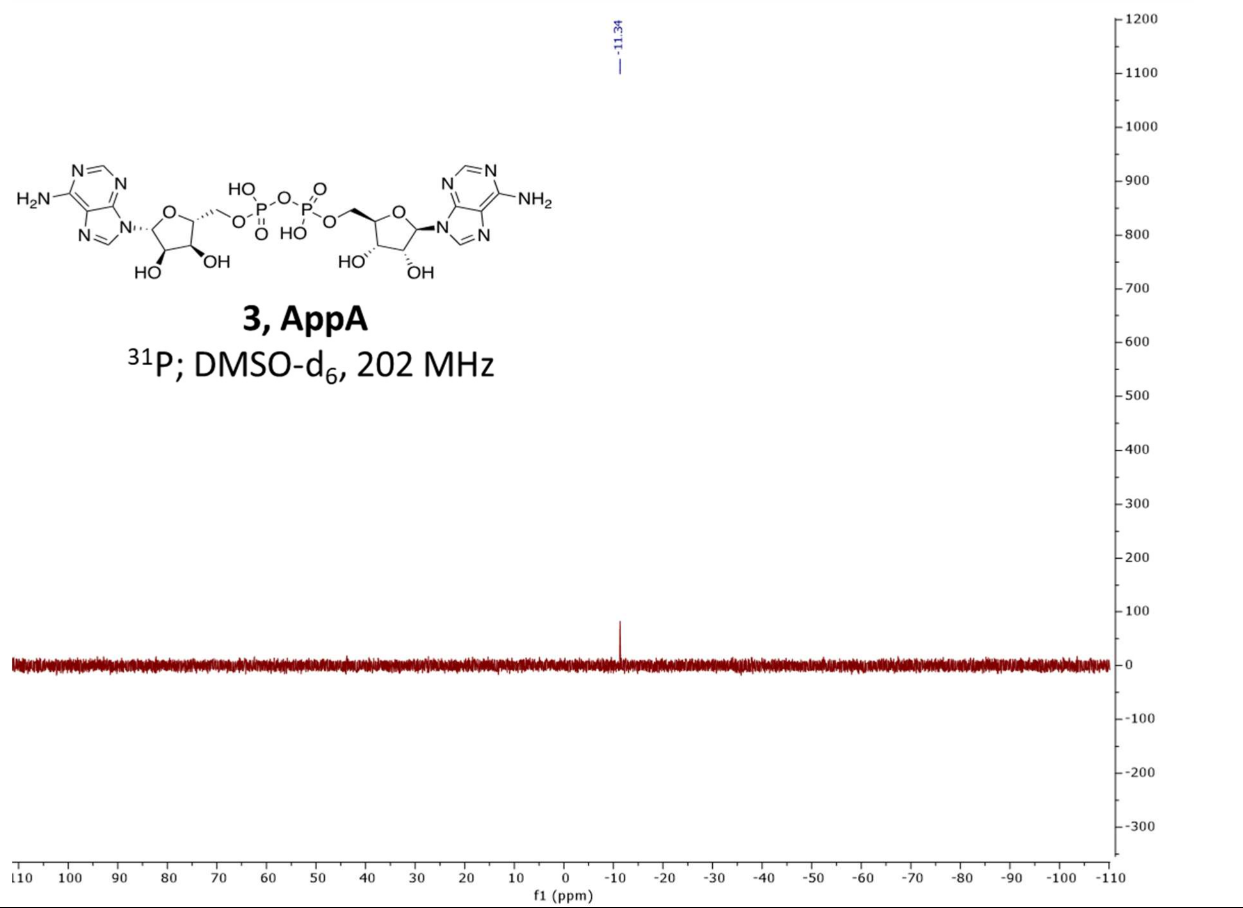


Figure S2. $^{31}$P-NMR of AppA synthetic standard

| | **Neutral Loss Mass (Da)** | **Product Ion $[M+H]^+$ (*m/z*)** |
|---|---|---|
| Deoxyribose | 116.0474 | --- |
| Guanine | 151.0494 | 152.0567 |
| Adenine | 135.0545 | 136.0617 |
| Thymine | 126.0429 | 127.0502 |
| Cytosine | 111.0433 | 112.0506 |
| Uracil | 112.0273 | 113.0346 |
| Guanine+$H_2O$ | 169.0600 | --- |
| Adenine+$H_2O$ | 153.0651 | --- |
| Ribose | 132.0423 | --- |
| 2 Deoxyribose | 232.0948 | --- |
| 2 Ribose | 264.0846 | --- |
| Ribityl | 134.0579 | --- |

Table S1. List of neutral losses and nucleobases monitored in the $MS^2$ to trigger $MS^3$

| *m/z* | *m/z* | *m/z* |
|---|---|---|
| 229.1431 | 397.1348 | 446.2040 |
| 241.1429 | 398.0773 | 447.126 |
| 258.1067 | 402.1768 | 451.0994 |
| 269.1378 | 408.1158 | 460.1834 |
| 364.1134 | 409.136 | 461.2026 |
| 371.1204 | 413.1219 | 462.1807 |
| 376.1619 | 414.177 | 469.1684 |
| 377.1453 | 415.1233 | 497.1834 |
| 378.1415 | 418.1726 | 507.2054 |
| 378.1493 | 419.118 | 539.1972 |
| 379.1279 | 421.1171 | 541.1926 |
| 379.1254 | 421.136 | 575.1736 |
| 381.1412 | 423.1523 | 585.2023 |
| **383.1529** | 429.1172 | 753.2837 |
| 387.1159 | 429.2337 | 799.2606 |
| 388.1614 | 430.1352 | 799.2884 |
| 388.1245 | 432.1873 | 1129.4291 |
| 391.1254 | 435.1519 | 1165.4001 |
| 393.1405 | 444.2239 | |

Table S2. List of m/z values used in the inclusion list for targeted DDA-CNL/$MS^3$ at high HCD. The *m/z* 383.1529 is for $^{13}C_4$,$^{15}N_2$-riboflavin.

| Measured Mass (Da) | Retention Time (min) | Adenine | Guanine | Cytosine | Uracil | Max. Area (n x $10^8$) | Charge State |
|---|---|---|---|---|---|---|---|
| 634.0946 | 9.4 | CC | | | | 8.1 | 1 |
| 652.1051 | 13.9 | CC | | CC | | 4.8 | 1 |
| 347.0635 | 13.9 | CC | | CC, NL | | 3.5 | 1 |
| 668.0993 | 13.5 | | CC | CC | | 3.1 | 1 |
| 653.0893 | 15 | CC | | | CC | 2.9 | 1 |
| 650.0893 | 13.8 | CC | | CC | | 2.8 | 1 |
| 596.1502 | 12.7 | | | CC | | 1.4 | 1 |
| 674.087 | 15.7 | CC, NL,* | | | | 0.95 | 1 |
| 612.1452 | 13.9 | CC, NL | | | | 0.92 | 1 |
| 676.1163 | 15.6 | CC, NL | CC, NL,** | CC | | 0.87 | 1 |
| 588.1341 | 14.7 | CC, NL | | | | 0.82 | 1 |
| 313.085 | 15.1 | CC, NL | | | | 0.77 | 1 |
| 963.148 | 14.4 | | CC, NL | CC | | 0.77 | 1 |
| 313.085 | 10.6 | CC | | | | 0.77 | 1 |
| 963.148 | 14.8 | CC, NL | | CC | | 0.75 | 2 |
| 329.053 | 13.5 | CC | | | | 0.59 | 1 |
| 692.1114 | 14.8 | CC | CC | | | 0.59 | 1 |
| 528.3025 | 13.3 | CC | | | | 0.59 | 2 |
| 981.1581 | 15.4 | CC | | CC | | 0.55 | 2 |
| 939.1364 | 13.7 | CC | | CC | | 0.48 | 2 |
| 658.1058 | 14.3 | CC | | | | 0.43 | 1 |
| 573.1229 | 15.9 | CC | | | CC | 0.43 | 1 |
| 348.1403 | 14.1 | CC | | CC | | 0.40 | 1 |
| 674.1007 | 14.5 | CC, NL | CC, NL | | | 0.35 | 1 |
| 696.0698 | 14 | CC | | CC | | 0.33 | 2 |
| 997.1535 | 15.6 | CC | CC | CC | | 0.28 | 2 |
| 979.1430 | 15.1 | CC | CC | CC | | 0.28 | 2 |
| 985.1300 | 14.7 | CC | | CC | | 0.27 | 2 |
| 1268.189 | 15.2 | CC | | CC | | 0.23 | 2 |
| 1292.2002 | 15.2 | CC | | CC | | 0.23 | 2 |
| 925.2038 | 16 | CC,* | | | | 0.21 | 2 |
| 961.1186 | 13.7 | CC | | CC | | 0.13 | 2 |
| 1244.1783 | 14.2 | CC | | CC | | 0.13 | 2 |

Table S3. Compounds identified in the adenine-based cluster with CC indicating observation via the Compound Class node as a protonated nucleobase and NL indicating observation of the neutral loss of one of the nucleobases. The primary charge state observed is indicated as either “1” or “2” indicating detection as $[M+1H]^{+1}$ and $[M+2H]^{+2}$, respectively. The * and ** symbols indicate observation of the neutral loss of adenosine and guanosine, respectively.

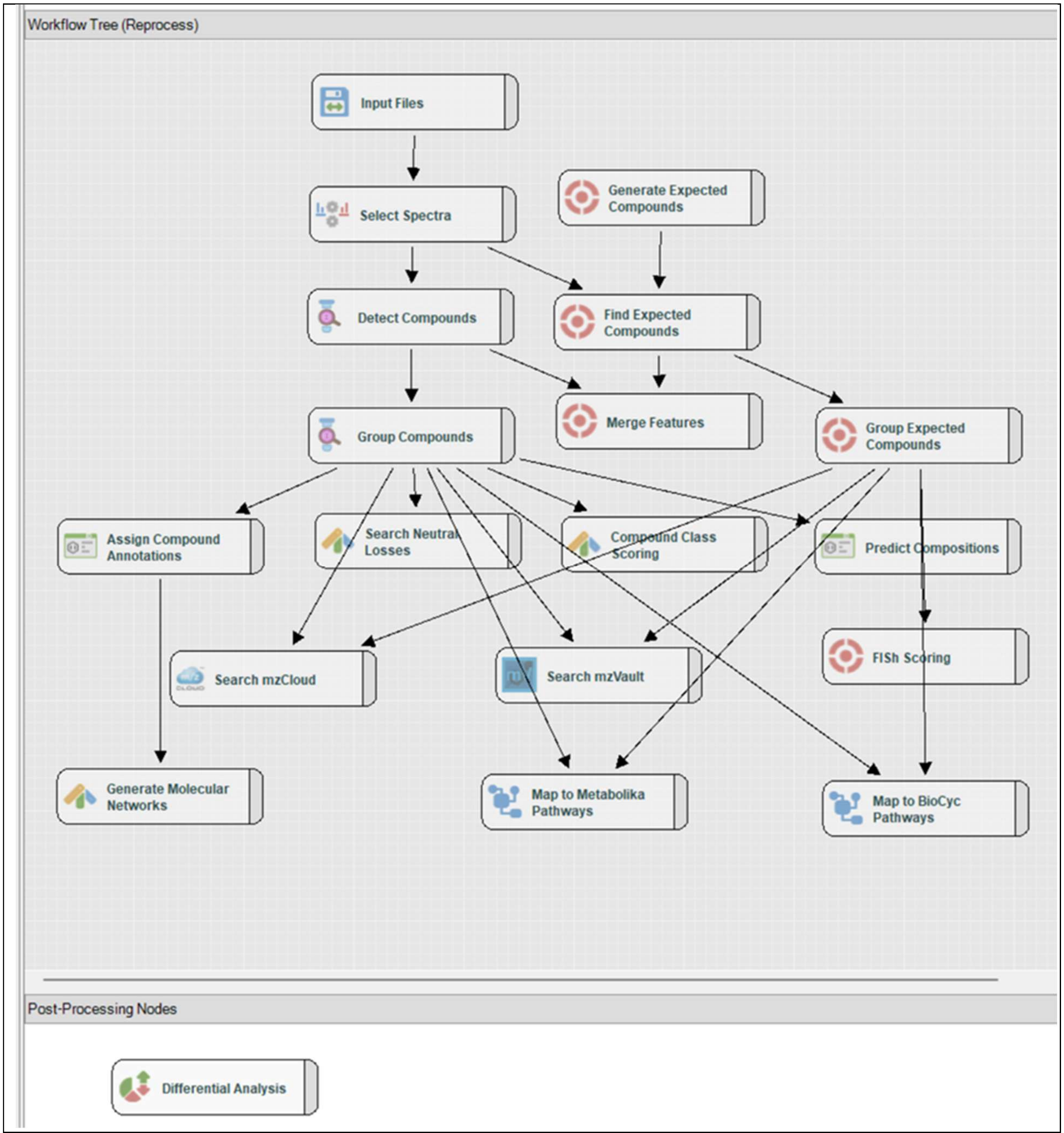


Scheme S2. Compound Discoverer workflow

**Compound Discoverer Workflow Parameters:**

Search name:
RibitylAdeninePhosphate_RiboflavinMetabolism_FullMetabolika_BioCyc_CL02_136_2
Search description: Untargeted food research ID workflow without statistics: Detect and identify unknown compounds.
- Performs retention time alignment, unknown compound detection, and compound grouping across all samples. Predicts elemental compositions for all compounds, and hides chemical background (using Blank samples). Identifies compounds using mzCloud (ddMS2 and/or DIA), ChemSpider (exact mass or formula) and local database searches against Mass Lists (exact mass with or without RT). Performs spectral similarity search against mzCloud for compounds with ddMS2. Applies mzLogic to rank order structure candidates from ChemSpider and mass list matches. And applies spectral distance scoring to ChemSpider and mass list matches.

Search date: 2/3/2025 5:30:21 PM
Created with Discoverer version: 3.3.3.200

[Input Files (0)]
    -->Select Spectra (36)

    [Select Spectra (36)]
        -->Find Expected Compounds (53)
        -->Detect Compounds (48)

        [Detect Compounds (48)]
            -->Group Compounds (47)
            -->Merge Features (55)

        [Generate Expected Compounds (54)]
            -->Find Expected Compounds (53)

            [Find Expected Compounds (53)]
                -->Merge Features (55)
                -->Group Expected Compounds (56)

            [Group Compounds (47)]
                -->Assign Compound Annotations (46)
                -->Compound Class Scoring (45)
                -->Search Neutral Losses (44)
                -->Search mzVault (50)
                -->Map to Metabolika Pathways (58)
                -->Map to BioCyc Pathways (59)
                -->Search mzCloud (49)
                -->Predict Compositions (26)

[Group Expected Compounds (56)]
-->FISh Scoring (57)
-->Search mzVault (50)
-->Map to Metabolika Pathways (58)
-->Map to BioCyc Pathways (59)
-->Search mzCloud (49)

[Assign Compound Annotations (46)]
-->Generate Molecular Networks (52)

[Merge Features (55)]

[FISh Scoring (57)]

[Search mzVault (50)]

[Map to Metabolika Pathways (58)]

[Map to BioCyc Pathways (59)]

[Search mzCloud (49)]

[Generate Molecular Networks (52)]

[Compound Class Scoring (45)]

[Search Neutral Losses (44)]

[Predict Compositions (26)]

[Differential Analysis (51)]

----------------------------------------------------------------
Processing node 0: Input Files
----------------------------------------------------------------
Input Data:
- File Name(s) (Hidden):
C:\Users\villa001\Desktop\Ribityl Phosphate Data\Untargeted Material for Ribityl Phosphate Paper\August 2022\CL02_136_CD1d_Batch1_Untargeted.raw
C:\Users\villa001\Desktop\Ribityl Phosphate Data\Untargeted Material for Ribityl Phosphate Paper\August 2022\CL02_136_CD1d_Batch2_Untargeted.raw
C:\Users\villa001\Desktop\Ribityl Phosphate Data\Untargeted Material for Ribityl Phosphate Paper\August 2022\CL02_136_MR1_Batch1_Untargeted.raw

C:\Users\villa001\Desktop\Ribityl Phosphate Data\Untargeted Material for Ribityl Phosphate Paper\August 2022\CL02_136_MR1_Batch2_Untargeted.raw

-----------------------------------------------------------------
Processing node 36: Select Spectra
-----------------------------------------------------------------
1. Spectrum Properties Filter:
- Lower RT Limit:  5
- Upper RT Limit:  50
- First Scan:  0
- Last Scan:  0
- Ignore Specified Scans:  (not specified)
- Total Intensity Threshold:  0
- Minimum Peak Count:  1

1.1 Spectrum Properties Filter for DDA Spectra:
- Lowest Charge State:  0
- Highest Charge State:  0
- Min. Precursor Mass:  0 Da
- Max. Precursor Mass:  5000 Da

2. Scan Event Filters:
- Mass Analyzer:  (not specified)
- MS Order:  Any
- Activation Type:  (not specified)
- Acquisition Type:  (not specified)
- Min. Collision Energy:  0
- Max. Collision Energy:  1000
- Scan Type:  Any
- Polarity Mode:  Is +
- MS1 Mass Range:  (not specified)
- FAIMS CV:  (not specified)

3. Peak Filters:
- S/N Threshold (FT-only):  1.5

4. Replacements for Unrecognized Properties:
- Unrecognized Charge Replacements:  1
- Unrecognized Mass Analyzer Replacements:  ITMS
- Unrecognized MS Order Replacements:  MS2
- Unrecognized Activation Type Replacements:  CID
- Unrecognized Polarity Replacements:  +
- Unrecognized MS Resolution@200 Replacements:  60000
- Unrecognized MSn Resolution@200 Replacements:  30000

6. General Settings:
- Precursor Selection:  Use MS(n - 1) Precursor
- Use Isotope Pattern in Precursor Reevaluation:  True
- Provide Profile Spectra:  Automatic
- Spectra to Store:  All
- Store Chromatograms:  False

----------------------------------------------------------------

Processing node 53: Find Expected Compounds

----------------------------------------------------------------

1. General Settings:
- Mass Tolerance:  5 ppm
- Intensity Tolerance [%]:  30
- Intensity Threshold [%]:  0.1
- Min. # Isotopes:  2
- Use Most Intense Isotope Only:  True
- Min. Peak Intensity:  1000
- Average Peak Width [min]:  0
- Precursor Mass Tolerance:  0.025 Da

2. Peak Detection:
- Chromatographic S/N Threshold:  1.5
- Remove Baseline:  False
- Gap Ratio Threshold:  0.35
- Max. Peak Width [min]:  1
- Min. Relative Valley Depth:  0.1
- Min. # Scans per Peak:  5

----------------------------------------------------------------

Processing node 55: Merge Features

----------------------------------------------------------------

1. Peak Consolidation:
- Mass Tolerance:  5 ppm
- RT Tolerance [min]:  0.05

----------------------------------------------------------------

Processing node 56: Group Expected Compounds

----------------------------------------------------------------

1. General Settings:
- RT Tolerance [min]:  0.2
- Minimum Valley [%]:  10
- Align Peaks:  False
- Preferred Ions:  [M+H]+1; [M-H]-1

- Area Integration:  Most Common Ion

2. Peak Rating Contributions:
- Area Contribution:  3
- CV Contribution:  10
- FWHM to Base Contribution:  5
- Jaggedness Contribution:  5
- Modality Contribution:  5
- Zig-Zag Index Contribution:  5

3. Peak Rating Filter:
- Peak Rating Threshold:  0
- Number of Files:  0

----------------------------------------------------------------
Processing node 57: FISh Scoring
----------------------------------------------------------------
1. General Settings:
- Annotate Full Tree:  True
- Match Transformations:  True
- S/N Threshold:  3
- High Acc. Mass Tolerance:  2.5 mmu
- Low Acc. Mass Tolerance:  0.5 Da

2. Fragment Prediction Settings:
- Use General Rules:  True
- Use Libraries:  True
- Max. Depth:  5
- Aromatic Cleavage:  True
- Min. Fragment m/z:  50

----------------------------------------------------------------
Processing node 50: Search mzVault
----------------------------------------------------------------
1. Search Settings:
- mzVault Library:  NIST 2020 MSMS HR.db
- Max. # Results:  10
- Match Factor Threshold:  50
- Search Algorithm:  HighChem HighRes
- Match Analyzer Type:  True
- IT Fragment Mass Tolerance:  0.4 Da
- FT Fragment Mass Tolerance:  10 ppm
- Use Retention Time:  False
- Precursor Mass Tolerance:  10 ppm

- Apply Intensity Threshold:  True
- Match Ionization Method:  False
- Ion Activation Energy Tolerance:  30
- Match Ion Activation Energy:  Any
- Match Ion Activation Type:  False
- Compound Classes:  All
- Remove Precursor Ion:  True
- RT Tolerance [min]:  2

----------------------------------------------------------------
Processing node 58: Map to Metabolika Pathways
----------------------------------------------------------------
1. Search Settings:
- Metabolika Pathways:  (3R)-linalool biosynthesis.metabolika|2-nitrobenzoate degradation I.metabolika|2-oxobutanoate degradation I.metabolika|3-phenylpropanoate and 3-(3-hydroxyphenyl)propanoate degradation.metabolika|3-phenylpropanoate degradation.metabolika|Acetyl-CoA fermentation to butanoate II.metabolika|Adenosylcobalamin biosynthesis I (anaerobic).metabolika|Adenosylcobalamin biosynthesis II (aerobic).metabolika|Allantoin degradation IV (anaerobic).metabolika|Allantoin degradation to glyoxylate I.metabolika|Allantoin degradation to glyoxylate II.metabolika|Allantoin degradation to glyoxylate III.metabolika|Ammonia assimilation cycle I.metabolika|Ammonia assimilation cycle III.metabolika|Ammonia oxidation IV (autotrophic ammonia oxidizers).metabolika|Anaerobic aromatic compound degradation (Thauera aromatica).metabolika|Anaerobic energy metabolism (invertebrates, mitochondrial).metabolika|Arachidonate biosynthesis III (6-desaturase, mammals).metabolika|Archaetidylinositol biosynthesis.metabolika|Archaetidylserine and archaetidylethanolamine biosynthesis.metabolika|Arginine, ornithine and proline interconversion.metabolika|Aromatic compounds degradation via ss-ketoadipate.metabolika|Aspartate superpathway.metabolika|B-carotene biosynthesis (engineered).metabolika|Bacillibactin biosynthesis.metabolika|Benzoate biosynthesis I (CoA-dependent, ss-oxidative).metabolika|Benzoate biosynthesis III (CoA-dependent, non-ss-oxidative).metabolika|Benzoate fermentation (to acetate and cyclohexane carboxylate).metabolika|Biotin biosynthesis I.metabolika|Biotin biosynthesis II.metabolika|Bitter acids biosynthesis.metabolika|Caffeine degradation IV (bacteria, via demethylation and oxidation).metabolika|Cardiolipin and phosphatidylethanolamine biosynthesis (Xanthomonas).metabolika|Catechol degradation I (meta-cleavage pathway).metabolika|Catechol degradation II (meta-cleavage pathway).metabolika|Catechol degradation III (ortho-cleavage pathway).metabolika|Cellulose and hemicellulose degradation (cellulolosome).metabolika|Chitin biosynthesis.metabolika|Cholesterol biosynthesis I.metabolika|Cholesterol biosynthesis II (via 24,25-dihydrolanosterol).metabolika|Cholesterol biosynthesis III (via desmosterol).metabolika|Choline degradation IV.metabolika|Choline-O-sulfate degradation.metabolika|Chondroitin sulfate biosynthesis.metabolika|Chorismate biosynthesis I.metabolika|Chorismate biosynthesis II (archaea).metabolika|Colanic acid building blocks biosynthesis.metabolika|Crotonate fermentation (to acetate and cyclohexane carboxylate).metabolika|Curcuminoid biosynthesis.metabolika|D-serine

metabolism.metabolika|Dermatan sulfate biosynthesis.metabolika|Enterobacterial common antigen biosynthesis.metabolika|Enterobactin biosynthesis.metabolika|G-butyrobetaine degradation.metabolika|GABA shunt.metabolika|Gamma-glutamyl cycle.metabolika|Gluconeogenesis II (Methanobacterium thermoautotrophicum).metabolika|Glycerol and glycerophosphodiester degradation.metabolika|Glycerol degradation to butanol.metabolika|Glycine biosynthesis II.metabolika|Heparan sulfate biosynthesis.metabolika|Hexitol fermentation to lactate, formate, ethanol and acetate.metabolika|Homolactic fermentation.metabolika|Hyperxanthone E biosynthesis.metabolika|Icosapentaenoate biosynthesis III (fungi).metabolika|Icosapentaenoate biosynthesis IV (bacteria).metabolika|Isoprene biosynthesis I.metabolika|Kanamycin biosynthesis.metabolika|Kauralexin biosynthesis.metabolika|Kdo transfer to lipid IVA III (Chlamydia).metabolika|Ketogluconate metabolism.metabolika|L-alanine fermentation to propanoate and acetate.metabolika|L-arginine biosynthesis I (via L-ornithine).metabolika|L-arginine degradation V (arginine deiminase pathway).metabolika|L-ascorbate biosynthesis V.metabolika|L-cysteine biosynthesis IV (from L-methionine).metabolika|L-cysteine biosynthesis IV (fungi).metabolika|L-glutamate and L-glutamine biosynthesis.metabolika|L-glutamate degradation IX (via 4-aminobutanoate).metabolika|L-glutamate degradation VII (to butanoate).metabolika|L-glutamate degradation VIII (to propanoate).metabolika|L-homoserine and L-methionine biosynthesis.metabolika|L-methionine biosynthesis III.metabolika|L-methionine salvage cycle I (bacteria and plants).metabolika|L-methionine salvage cycle II (plants).metabolika|L-methionine salvage cycle III.metabolika|L-tryptophan degradation III (eukaryotic).metabolika|L-tryptophan degradation IX.metabolika|L-tryptophan degradation XI (mammalian, via kynurenine).metabolika|L-tryptophan degradation XII (Geobacillus).metabolika|L-tyrosine degradation IV (to 4-methylphenol).metabolika|Mandelate degradation to acetyl-CoA.metabolika|Meta cleavage pathway of aromatic compounds.metabolika|Methanobacterium thermoautotrophicum biosynthetic metabolism.metabolika|Methanol and methylamine oxidation to formaldehyde.metabolika|Methanol oxidation to carbon dioxide.metabolika|Methylglyoxal degradation IV.metabolika|MRNA capping II.metabolika|Myo-, chiro- and scillo-inositol degradation.metabolika|N-acetylglucosamine degradation II.metabolika|NAD biosynthesis II (from tryptophan).metabolika|NAD salvage pathway III.metabolika|Naphthalene degradation to acetyl-CoA.metabolika|Nitrifier denitrification.metabolika|Novobiocin biosynthesis.metabolika|O-antigen building blocks biosynthesis (E. coli).metabolika|Oxygenic photosynthesis.metabolika|P-cumate degradation.metabolika|P-cymene degradation.metabolika|Pentose phosphate pathway.metabolika|Peptidoglycan biosynthesis I (meso-diaminopimelate containing).metabolika|Peptidoglycan biosynthesis II (staphylococci).metabolika|Peptidoglycan biosynthesis III (mycobacteria).metabolika|Peptidoglycan biosynthesis IV (Enterococcus faecium).metabolika|Peptidoglycan biosynthesis V (ss-lactam resistance).metabolika|Phosphatidylglycerol biosynthesis I (plastidic).metabolika|Phosphatidylglycerol biosynthesis II (non-plastidic).metabolika|Plant sterol biosynthesis.metabolika|Polyisoprenoid biosynthesis (E. coli).metabolika|Purine nucleotides degradation I (plants).metabolika|Purine nucleotides degradation II (aerobic).metabolika|Pyrimidine nucleobases salvage II.metabolika|Pyruvate fermentation to acetate and alanine.metabolika|Pyruvate fermentation to acetate and lactate

I.metabolika|Pyruvate fermentation to acetate and lactate II.metabolika|Pyruvate fermentation to acetate I.metabolika|Pyruvate fermentation to acetate III.metabolika|Pyruvate fermentation to acetate IV.metabolika|Pyruvate fermentation to acetate V.metabolika|Pyruvate fermentation to acetate VI.metabolika|Pyruvate fermentation to acetate VII.metabolika|Reactive oxygen species degradation.metabolika|S-adenosyl-L-methionine cycle I.metabolika|Salicylate glucosides biosynthesis I.metabolika|Sphingolipid biosynthesis (mammals).metabolika|Sucrose biosynthesis I (from photosynthesis).metabolika|Sulfate reduction I (assimilatory).metabolika|Superpathway avenacin A biosynthesis.metabolika|Superpathway NADNADP - NADHNADPH interconversion (yeast).metabolika|Superpathway of (Kdo)2-lipid A biosynthesis.metabolika|Superpathway of (R,R)-butanediol biosynthesis.metabolika|Superpathway of 1D-myo-inositol hexakisphosphate biosynthesis (plants).metabolika|Superpathway of 2,3-butanediol biosynthesis.metabolika|Superpathway of 4-aminobutanoate degradation.metabolika|Superpathway of 4-hydroxybenzoate biosynthesis (yeast).metabolika|Superpathway of 5-aminoimidazole ribonucleotide biosynthesis.metabolika|Superpathway of acetate utilization and formation.metabolika|Superpathway of acetyl-CoA biosynthesis.metabolika|Superpathway of acrylonitrile degradation.metabolika|Superpathway of adenosine nucleotides de novo biosynthesis I.metabolika|Superpathway of adenosine nucleotides de novo biosynthesis II.metabolika|Superpathway of aerobic toluene degradation.metabolika|Superpathway of aflatoxin biosynthesis.metabolika|Superpathway of allantoin degradation in plants.metabolika|Superpathway of allantoin degradation in yeast.metabolika|Superpathway of Allium flavor precursors.metabolika|Superpathway of ammonia assimilation (plants).metabolika|Superpathway of anaerobic energy metabolism (invertebrates).metabolika|Superpathway of anaerobic sucrose degradation.metabolika|Superpathway of anthocyanin biosynthesis (from cyanidin and cyanidin 3-O-glucoside).metabolika|Superpathway of anthocyanin biosynthesis (from delphinidin 3-O-glucoside).metabolika|Superpathway of anthocyanin biosynthesis (from pelargonidin 3-O-glucoside).metabolika|Superpathway of arginine and polyamine biosynthesis.metabolika|Superpathway of aromatic amino acid biosynthesis.metabolika|Superpathway of aromatic compound degradation via 2-oxopent-4-enoate.metabolika|Superpathway of aromatic compound degradation via 3-oxoadipate.metabolika|Superpathway of atrazine degradation.metabolika|Superpathway of bacteriochlorophyll a biosynthesis.metabolika|Superpathway of benzoxazinoid glucosides biosynthesis.metabolika|Superpathway of betalain biosynthesis.metabolika|Superpathway of branched chain amino acid biosynthesis.metabolika|Superpathway of butirocin biosynthesis.metabolika|Superpathway of C1 compounds oxidation to CO2.metabolika|Superpathway of C28 brassinosteroid biosynthesis.metabolika|Superpathway of candicidin biosynthesis.metabolika|Superpathway of carotenoid biosynthesis.metabolika|Superpathway of CDP-glucose-derived O-antigen building blocks biosynthesis.metabolika|Superpathway of cholesterol biosynthesis.metabolika|Superpathway of cholesterol degradation I (cholesterol oxidase).metabolika|Superpathway of cholesterol degradation II (cholesterol dehydrogenase).metabolika|Superpathway of choline biosynthesis.metabolika|Superpathway of chorismate metabolism.metabolika|Superpathway of CMP-sialic acids biosynthesis.metabolika|Superpathway of coenzyme A biosynthesis

I.metabolika|Superpathway of coenzyme A biosynthesis II (plants).metabolika|Superpathway of coenzyme A biosynthesis III (mammals).metabolika|Superpathway of cytosolic glycolysis (plants), pyruvate dehydrogenase and TCA cycle.metabolika|Superpathway of D-glucarate and D-galactarate degradation.metabolika|Superpathway of D-myo-inositol (1,4,5)-trisphosphate metabolism.metabolika|Superpathway of demethylmenaquinol-6 biosynthesis I.metabolika|Superpathway of demethylmenaquinol-6 biosynthesis II.metabolika|Superpathway of demethylmenaquinol-8 biosynthesis.metabolika|Superpathway of demethylmenaquinol-9 biosynthesis.metabolika|Superpathway of dimethylsulfone degradation.metabolika|Superpathway of dimethylsulfoniopropanoate degradation.metabolika|Superpathway of diterpene resin acids biosynthesis.metabolika|Superpathway of dTDP-glucose-derived antibiotic building blocks biosynthesis.metabolika|Superpathway of dTDP-glucose-derived O-antigen building blocks biosynthesis.metabolika|Superpathway of ergosterol biosynthesis I.metabolika|Superpathway of ergosterol biosynthesis II.metabolika|Superpathway of ergotamine biosynthesis.metabolika|Superpathway of erythromycin biosynthesis (without sugar biosynthesis).metabolika|Superpathway of erythromycin biosynthesis.metabolika|Superpathway of fatty acid biosynthesis I (E. coli).metabolika|Superpathway of fatty acid biosynthesis II (plant).metabolika|Superpathway of fatty acid biosynthesis initiation (E. coli).metabolika|Superpathway of fatty acids biosynthesis (E. coli).metabolika|Superpathway of fermentation (Chlamydomonas reinhardtii).metabolika|Superpathway of flavones and derivatives biosynthesis .metabolika|Superpathway of formononetin derivative biosynthesis.metabolika|Superpathway of fucose and rhamnose degradation.metabolika|Superpathway of fumitremorgin biosynthesis.metabolika|Superpathway of GDP-mannose-derived O-antigen building blocks biosynthesis.metabolika|Superpathway of geranylgeranyl diphosphate biosynthesis II (via MEP).metabolika|Superpathway of geranylgeranyldiphosphate biosynthesis I (via mevalonate).metabolika|Superpathway of gibberellin biosynthesis.metabolika|Superpathway of gibberellin GA12 biosynthesis.metabolika|Superpathway of glucose and xylose degradation.metabolika|Superpathway of glycerol degradation to 1,3-propanediol.metabolika|Superpathway of glycol metabolism and degradation.metabolika|Superpathway of glycolysis and Entner-Doudoroff.metabolika|Superpathway of glycolysis, pyruvate dehydrogenase, TCA, and glyoxylate bypass.metabolika|Superpathway of glyoxylate bypass and TCA.metabolika|Superpathway of glyoxylate cycle and fatty acid degradation.metabolika|Superpathway of guanine and guanosine salvage.metabolika|Superpathway of guanosine nucleotides degradation (plants).metabolika|Superpathway of guanosine nucleotides de novo biosynthesis I.metabolika|Superpathway of guanosine nucleotides de novo biosynthesis II.metabolika|Superpathway of heme biosynthesis from glutamate.metabolika|Superpathway of heme biosynthesis from glycine.metabolika|Superpathway of heme biosynthesis from uroporphyrinogen-III.metabolika|Superpathway of hexitol degradation (bacteria).metabolika|Superpathway of hexuronide and hexuronate degradation.metabolika|Superpathway of histidine, purine, and pyrimidine biosynthesis.metabolika|Superpathway of hydrogen production.metabolika|Superpathway of hydrolyzable tannin biosynthesis.metabolika|Superpathway of hyoscyamine and scopolamine biosynthesis.metabolika|Superpathway of indole-3-acetate conjugate biosynthesis.metabolika|Superpathway of inositol phosphate compounds.metabolika|Superpathway of isoflavonoids (via naringenin).metabolika|Superpathway of jasmonoyl-amino acid conjugates biosynthesis.metabolika|Superpathway of L-alanine biosynthesis.metabolika|Superpathway of L-arginine and L-ornithine degradation.metabolika|Superpathway of L-arginine, putrescine, and 4-aminobutanoate degradation.metabolika|Superpathway of L-asparagine biosynthesis.metabolika|Superpathway of L-aspartate and L-asparagine biosynthesis.metabolika|Superpathway of L-citrulline metabolism.metabolika|Superpathway of L-cysteine biosynthesis (mammalian).metabolika|Superpathway of L-isoleucine biosynthesis I.metabolika|Superpathway of L-lysine degradation.metabolika|Superpathway of L-lysine, L-threonine and L-methionine biosynthesis I.metabolika|Superpathway of L-lysine, L-threonine and L-methionine biosynthesis II.metabolika|Superpathway of L-methionine biosynthesis (by sulfhydrylation).metabolika|Superpathway of L-methionine biosynthesis (transsulfuration).metabolika|Superpathway of L-methionine salvage and degradation.metabolika|Superpathway of L-phenylalanine and L-tyrosine biosynthesis.metabolika|Superpathway of L-phenylalanine biosynthesis.metabolika|Superpathway of L-serine and glycine biosynthesis I.metabolika|Superpathway of L-threonine biosynthesis.metabolika|Superpathway of L-threonine metabolism.metabolika|Superpathway of L-tryptophan biosynthesis.metabolika|Superpathway of L-tyrosine biosynthesis.metabolika|Superpathway of linalool biosynthesis.metabolika|Superpathway of linamarin and lotaustralin biosynthesis.metabolika|Superpathway of lipopolysaccharide biosynthesis.metabolika|Superpathway of lipoxygenase.metabolika|Superpathway of megalomicin A biosynthesis.metabolika|Superpathway of melatonin degradation.metabolika|Superpathway of menaquinol-10 biosynthesis.metabolika|Superpathway of menaquinol-11 biosynthesis.metabolika|Superpathway of menaquinol-12 biosynthesis.metabolika|Superpathway of menaquinol-13 biosynthesis.metabolika|Superpathway of menaquinol-6 biosynthesis I.metabolika|Superpathway of menaquinol-7 biosynthesis.metabolika|Superpathway of menaquinol-8 biosynthesis I.metabolika|Superpathway of menaquinol-8 biosynthesis II.metabolika|Superpathway of menaquinol-9 biosynthesis.metabolika|Superpathway of methanogenesis.metabolika|Superpathway of methylglyoxal degradation.metabolika|Superpathway of microbial D-galacturonate and D-glucuronate degradation.metabolika|Superpathway of mycolyl-arabinogalactan-peptidoglycan complex biosynthesis.metabolika|Superpathway of NAD biosynthesis in eukaryotes.metabolika|Superpathway of neomycin biosynthesis.metabolika|Superpathway of nicotinate degradation.metabolika|Superpathway of nicotine biosynthesis.metabolika|Superpathway of oleoresin turpentine biosynthesis.metabolika|Superpathway of ornithine degradation.metabolika|Superpathway of penicillin, cephalosporin and cephamycin biosynthesis.metabolika|Superpathway of pentose and pentitol degradation.metabolika|Superpathway of phenylethylamine degradation.metabolika|Superpathway of phosphatidylcholine biosynthesis.metabolika|Superpathway of phospholipid biosynthesis I (bacteria).metabolika|Superpathway of phospholipid biosynthesis II (plants).metabolika|Superpathway of photosynthetic hydrogen production.metabolika|Superpathway of phylloquinol biosynthesis.metabolika|Superpathway of plastoquinol biosynthesis.metabolika|Superpathway of polyamine biosynthesis I.metabolika|Superpathway of polyamine biosynthesis II.metabolika|Superpathway of polyamine biosynthesis III.metabolika|Superpathway of pterocarpan biosynthesis (via daidzein).metabolika|Superpathway of pterocarpan biosynthesis (via formononetin).metabolika|Superpathway of purine deoxyribonucleosides degradation.metabolika|Superpathway of purine nucleotide salvage.metabolika|Superpathway of purine nucleotides de novo biosynthesis I.metabolika|Superpathway of purine nucleotides de novo biosynthesis II.metabolika|Superpathway of purines degradation in plants.metabolika|Superpathway of pyridoxal 5'-phosphate biosynthesis and salvage.metabolika|Superpathway of pyrimidine deoxyribonucleoside salvage.metabolika|Superpathway of pyrimidine deoxyribonucleosides degradation.metabolika|Superpathway of pyrimidine deoxyribonucleotides de novo biosynthesis (E. coli).metabolika|Superpathway of pyrimidine deoxyribonucleotides de novo biosynthesis.metabolika|Superpathway of pyrimidine nucleobases salvage.metabolika|Superpathway of pyrimidine ribonucleosides degradation.metabolika|Superpathway of pyrimidine ribonucleosides salvage.metabolika|Superpathway of pyrimidine ribonucleotides de novo biosynthesis.metabolika|Superpathway of quinolone and alkylquinolone biosynthesis.metabolika|Superpathway of rifamycin B biosynthesis.metabolika|Superpathway of roquefortine, meleagrin and neoxaline biosynthesis.metabolika|Superpathway of rosmarinic acid biosynthesis.metabolika|Superpathway of salicylate degradation.metabolika|Superpathway of scopolin and esculin biosynthesis.metabolika|Superpathway of seleno-compound metabolism.metabolika|Superpathway of ss-D-glucuronide and D-glucuronate degradation.metabolika|Superpathway of stearidonate biosynthesis (cyanobacteria).metabolika|Superpathway of steroid hormone biosynthesis.metabolika|Superpathway of sulfate assimilation and cysteine biosynthesis.metabolika|Superpathway of sulfide oxidation (Acidithiobacillus ferrooxidans).metabolika|Superpathway of sulfide oxidation (phototrophic sulfur bacteria).metabolika|Superpathway of sulfide oxidation (Starkeya novella).metabolika|Superpathway of sulfolactate degradation.metabolika|Superpathway of sulfur amino acid biosynthesis (Saccharomyces cerevisiae).metabolika|Superpathway of sulfur metabolism (Desulfocapsa sulfoexigens).metabolika|Superpathway of sulfur oxidation (Acidianus ambivalens).metabolika|Superpathway of taurine degradation.metabolika|Superpathway of testosterone and androsterone degradation.metabolika|Superpathway of tetracycline and oxytetracycline biosynthesis.metabolika|Superpathway of tetrahydrofolate biosynthesis and salvage.metabolika|Superpathway of tetrahydrofolate biosynthesis.metabolika|Superpathway of tetrahydroxyxanthone biosynthesis.metabolika|Superpathway of tetrathionate reduction (Salmonella typhimurium).metabolika|Superpathway of the 3-hydroxypropanoate

cycle.metabolika|Superpathway of thiamine diphosphate biosynthesis I.metabolika|Superpathway of thiamine diphosphate biosynthesis II.metabolika|Superpathway of thiamine diphosphate biosynthesis III (eukaryotes).metabolika|Superpathway of thiosulfate metabolism (Desulfovibrio sulfodismutans).metabolika|Superpathway of trichothecene biosynthesis.metabolika|Superpathway of trimethylamine degradation.metabolika|Superpathway of ubiquinol-6 biosynthesis (eukaryotic).metabolika|Superpathway of ubiquinol-8 biosynthesis (prokaryotic).metabolika|Superpathway of UDP-glucose-derived O-antigen building blocks biosynthesis.metabolika|Superpathway of UDP-N-acetylglucosamine-derived O-antigen building blocks biosynthesis.metabolika|Superpathway of unsaturated fatty acids biosynthesis (E. coli).metabolika|Superpathway of vanillin and vanillate degradation.metabolika|Superpathway of Clostridium acetobutylicum acidogenic and solventogenic fermentation.metabolika|Superpathway of Clostridium acetobutylicum acidogenic fermentation.metabolika|Superpathway of Clostridium acetobutylicum solventogenic fermentation.metabolika|Superpathway of N-acetylglucosamine, N-acetylmannosamine and N-acetylneuraminate degradation.metabolika|Superpathway of N-acetylneuraminate degradation.metabolika|Superpathway of S-adenosyl-L-methionine biosynthesis.metabolika|Superpathway polymethylated quercetinquercetagetin glucoside biosynthesis (Chrysosplenium).metabolika|Superpathways of coenzyme A biosynthesis I.metabolika|Superpathways of coenzyme A biosynthesis III (mammals).metabolika|Syringate degradation.metabolika|Taxadiene biosynthesis (engineered).metabolika|Thiamine salvage II.metabolika|Toluene degradation I (aerobic) (via o-cresol).metabolika|Toluene degradation II (aerobic) (via 4-methylcatechol).metabolika|Toluene degradation III (aerobic) (via p-cresol).metabolika|Toluene degradation IV (aerobic) (via catechol).metabolika|Toluene degradation V (aerobic) (via toluene-cis-diol).metabolika|Toluene degradation VI (anaerobic).metabolika|Trans-lycopene biosynthesis I (bacteria).metabolika|UDP-D-xylose biosynthesis.metabolika|UDP-galactofuranose biosynthesis.metabolika|UDP-sugars interconversion.metabolika|Ureide biosynthesis.metabolika|Vibriobactin biosynthesis.metabolika|Wybutosine biosynthesis.metabolika
- Search Mode:  By Mass Only

2. By Mass Search Settings:
- Mass Tolerance:  5 ppm

3. By Formula Search Settings:
- Max. # of Predicted Compositions to be searched per Compound:  3

4. Display Settings:
- Max. # Pathways in 'Pathways' column:  20

----------------------------------------------------------------
Processing node 59: Map to BioCyc Pathways
----------------------------------------------------------------
1. Search Settings:
- BioCyc Database/organism to be searched:  Homo sapiens  (HUMAN)

- Search Mode:  By Mass Only

2. By Mass Search Settings:
- Mass Tolerance:  5 ppm

3. By Formula Search Settings:
- Max. # of Predicted Compositions to be searched per Compound:  3

4. Display Settings:
- Max. # Pathways in 'Pathways' column:  20

----------------------------------------------------------------
Processing node 49: Search mzCloud
----------------------------------------------------------------
1. General Settings:
- Compound Classes:  All
- Precursor Mass Tolerance:  10 ppm
- FT Fragment Mass Tolerance:  10 ppm
- IT Fragment Mass Tolerance:  0.4 Da
- Library:  Autoprocessed; Reference
- Post Processing:  Recalibrated
- Max. # Results:  10
- Annotate Matching Fragments:  False
- Search MSn Tree:  True

2. DDA Search:
- Identity Search:  HighChem HighRes
- Match Activation Type:  False
- Match Activation Energy:  Any
- Activation Energy Tolerance:  30
- Apply Intensity Threshold:  True
- Similarity Search:  Similarity Forward
- Match Factor Threshold:  60

3. DIA Search:
- Use DIA Scans for Search:  False
- Max. Isolation Width [Da]:  500
- Match Activation Type:  False
- Match Activation Energy:  Any
- Activation Energy Tolerance:  100
- Apply Intensity Threshold:  False
- Match Factor Threshold:  20

----------------------------------------------------------------

Processing node 48: Detect Compounds
----------------------------------------------------------------
1. General Settings:
- Mass Tolerance [ppm]:  10 ppm
- Min. Peak Intensity:  1000000
- Min. # Scans per Peak:  5
- Use Most Intense Isotope Only:  True
- Precursor Mass Tolerance:  10 ppm

2. Trace Detection:
- Max. Number of Gaps to Correct:  2
- Min. Number of Adjacent Non-Zeros:  2
- Trace Mass Update Strategy:  Weighted Mean

3. Peak Detection:
- Chromatographic S/N Threshold:  1.5
- Remove Baseline:  False
- Gap Ratio Threshold:  0.35
- Max. Peak Width [min]:  1
- Min. Relative Valley Depth:  0.1

4. Isotope Pattern Detection:
- Group Isotopes for:  (not specified)
- RT Tolerance [min]:  0
- Use Peak Quality for Isotope Grouping:  True
- Filter out Features with Bad Peaks Only:  True
- Zig-Zag Index Threshold:  0.2
- Jaggedness Threshold:  0.4
- Modality Threshold:  0.9
- Remove Potentially False Positive Isotopes:  True

5. Compound Assembly:
- Ions:  [M+2H]+2; [M+H]+1
- Base Ions:  [M+H]+1; [M-H]-1
- Remove Singlets:  True

6. AcquireX Settings:
- Detect Persistent Background Ions:  False

----------------------------------------------------------------
Processing node 47: Group Compounds
----------------------------------------------------------------
1. General Settings:
- Mass Tolerance:  10 ppm

- RT Tolerance [min]:  0.5
- Minimum Valley [%]:  10
- Align Peaks:  False
- Preferred Ions:  [M+H]+1
- Area Integration:  Most Common Ion

2. Peak Rating Contributions:
- Area Contribution:  3
- CV Contribution:  10
- FWHM to Base Contribution:  5
- Jaggedness Contribution:  5
- Modality Contribution:  5
- Zig-Zag Index Contribution:  5

3. Peak Rating Filter:
- Peak Rating Threshold:  0
- Number of Files:  0

----------------------------------------------------------------
Processing node 46: Assign Compound Annotations
----------------------------------------------------------------
1. General Settings:
- Mass Tolerance:  10 ppm

2. Data Sources:
- Data Source #1:  mzCloud Search
- Data Source #2:  mzVault Search
- Data Source #3:  Metabolika Search
- Data Source #4:  BioCyc Search
- Data Source #5:  ChemSpider Search
- Data Source #6:  Predicted Compositions
- Data Source #7:  (not specified)

3. Scoring Rules:
- Use mzLogic:  True
- Use Spectral Distance:  True
- SFit Threshold:  20
- SFit Range:  20

4. Reprocessing:
- Clear Names:  False

----------------------------------------------------------------
Processing node 52: Generate Molecular Networks

----------------------------------------------------------------
1. Spectral Similarity:
- Use Full MSn Tree:  True
- Match Mass Shift:  True
- Match Transformations:  False
- Variate Transformations:  False
- S/N Threshold:  10
- Mass Tolerance:  10 ppm
- Min. Fragment m/z:  50

2. Transformations:
- Phase I:  (not specified)
- Phase II:  (not specified)
- Others:  (not specified)
- Max. # Phase II:  0
- Max. # All Steps:  0

3. Applied View Filters:
- Require Transformation:  False
- Require MSn:  True
- Min. MSn Score:  50
- Min. MSn Coverage:  70
- Min. Fragments:  3

4. Applied Thresholds:
- Require Transformation:  False
- Require MSn:  False
- Min. MSn Score:  20
- Min. MSn Coverage:  20
- Min. Fragments:  0

----------------------------------------------------------------
Processing node 45: Compound Class Scoring
----------------------------------------------------------------
1. General Settings:
- Compound Classes:  Adenine.cLib|Guanine.cLib|Cytosine.cLib|Thymine.cLib|Uracil.cLib
- S/N Threshold:  10
- High Acc. Mass Tolerance:  10 ppm
- Low Acc. Mass Tolerance:  10 ppm
- Use Full MS Tree:  False
- Allow DIA Scoring:  False

----------------------------------------------------------------
Processing node 44: Search Neutral Losses

----------------------------------------------------------------
1. General Settings:
- Neutral Losses:
    Adenine (C5 H5 N5, 135.05)
    AdenineH2O (C5 H7 N5 O, 153.07)
    Adenosine (C10 H13 N5 O4, 267.10)
    Cytosine (C4 H5 N3 O, 111.04)
    Deoxyribose (C5 H8 O3, 116.05)
    Guanine (C5 H5 N5 O, 151.05)
    GuanineH2O (C5 H7 N5 O2, 169.06)
    Guanosine (C10 H13 N5 O5, 283.09)
    Ribityl (C5 H10 O4, 134.06)
    Ribose (C5 H8 O4, 132.04)
    Thymine (C5 H6 N2 O2, 126.04)
    Uracil (C4 H4 N2 O2, 112.03)
- High Acc. Mass Tolerance:  10 ppm
- Low Acc. Mass Tolerance:  10 ppm
- S/N Threshold:  30
- Use DIA Scans for Search:  False

----------------------------------------------------------------
Processing node 26: Predict Compositions
----------------------------------------------------------------
1. Prediction Settings:
- Mass Tolerance:  10 ppm
- Min. Element Counts:  C H
- Max. Element Counts:  C90 H190 N10 O18 P3 S2
- Min. RDBE:  0
- Max. RDBE:  40
- Min. H/C:  0.1
- Max. H/C:  3.5
- Max. # Candidates:  10
- Max. # Internal Candidates:  500

2. Pattern Matching:
- Intensity Tolerance [%]:  30
- Intensity Threshold [%]:  0.1
- S/N Threshold:  3
- Min. Spectral Fit [%]:  30
- Min. Pattern Cov. [%]:  80
- Use Dynamic Recalibration:  True

3. Fragments Matching:
- Use Fragments Matching:  True

- Mass Tolerance:  5 ppm
- S/N Threshold:  3

----------------------------------------------------------------
Processing node 54: Generate Expected Compounds
----------------------------------------------------------------
1. Compound Selection:
- Compounds:  Riboflavin (C17 H20 N4 O6)

2. Dealkylation:
- Apply Dealkylation:  True
- Apply Dearylation:  True
- Max. # Steps:  1
- Min. Mass [Da]:  200

3. Transformations:
- Phase I:
    Dehydration (H2 O -> )
    Desaturation (H2 -> )
    Hydration ( -> H2 O)
    Nitro Reduction (O2 -> H2)
    Oxidation ( -> O)
    Oxidative Deamination to Alcohol (H2 N -> H O)
    Oxidative Deamination to Ketone (H3 N -> O)
    Reduction ( -> H2)
    Thiourea to Urea (S -> O)
- Phase II:
    Acetylation (H -> C2 H3 O)
    Arginine Conjugation (H O -> C6 H13 N4 O2)
    Cysteine Conjugation 1 (H -> C3 H6 N O2 S)
    Cysteine Conjugation 2 ( -> C3 H7 N O2 S)
    Glucoside Conjugation (H -> C6 H11 O5)
    Glucuronide Conjugation (H -> C6 H9 O6)
    Glutamine Conjugation (H O -> C5 H9 N2 O3)
    Glycine Conjugation (H O -> C2 H4 N O2)
    GSH Conjugation 1 ( -> C10 H15 N3 O6 S)
    GSH Conjugation 2 ( -> C10 H17 N3 O6 S)
    Methylation (H -> C H3)
    Ornitine Conjugation (H O -> C5 H11 N2 O2)
    Palmitoyl Conjugation (H -> C16 H31 O)
    Stearyl Conjugation (H -> C18 H35 O)
    Sulfation (H -> H O3 S)
    Taurine Conjugation (H O -> C2 H6 N O3 S)
- Others:  (not specified)

- Max. # Phase II:  1
- Max. # All Steps:  4

4. Ionization:
- Ions:  [M+H]+1

----------------------------------------------------------------
Processing node 51: Differential Analysis
----------------------------------------------------------------
1. General Settings:
- Log10 Transform Values:  True
- Group Area Calculation:  Median
- Replicate Area Calculation:  Median

2. Peak Rating Contributions:
- Update Peak Rating:  True
- Area Contribution:  3
- CV Contribution:  10
- FWHM to Base Contribution:  5
- Jaggedness Contribution:  5
- Modality Contribution:  5
- Zig-Zag Index Contribution:  5

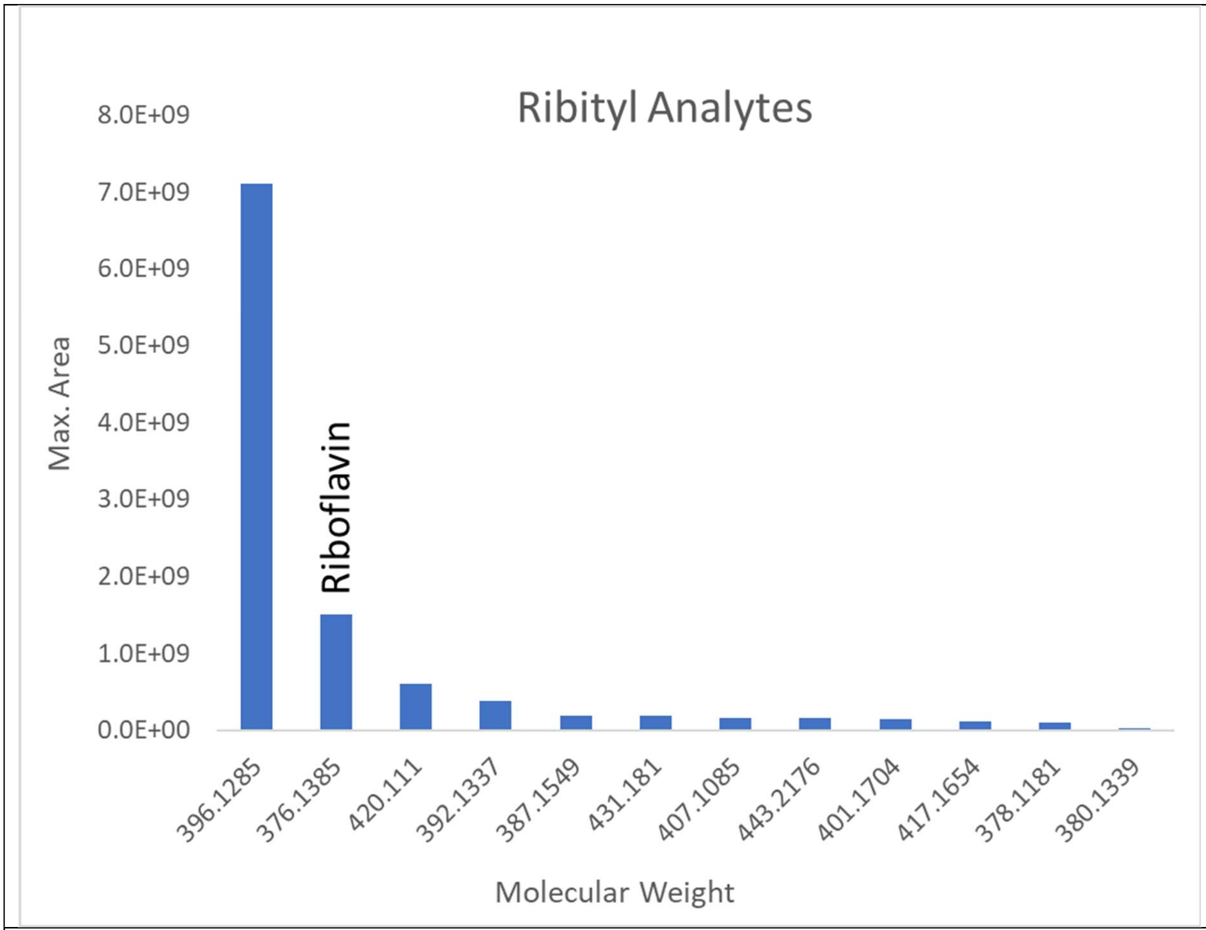


Figure S3. Extracted protonated ion chromatographic peak areas for ribityl analytes listed in Table 1.

| MS$^2$, *m/z* 397, HCD 55 | | | MS$^2$, *m/z* 397, HCD 55<br>MS$^3$ *m/z* 263, HCD 55 | |
|---|---|---|---|---|
| *m/z* | Relative Abundance (%) | | *m/z* | Relative Abundance (%) |
| 188.0812 | 100.0 | | 188.0813 | 100.0 |
| 259.0823 | 94.2 | | 214.0607 | 70.0 |
| 232.0712 | 59.0 | | 161.0704 | 64.7 |
| 69.0330 | 23.0 | | 186.0656 | 41.3 |
| 214.0607 | 15.1 | | 133.0753 | 40.0 |
| 234.0870 | 13.1 | | 146.0593 | 35.7 |
| 216.0762 | 10.7 | | 173.0578 | 32.9 |
| 161.0705 | 9.5 | | 158.0707 | 30.0 |
| 190.0970 | 8.0 | | 91.0536 | 20.2 |
| 81.0330 | 6.5 | | 160.0864 | 20.1 |
| 57.0331 | 6.5 | | 232.0714 | 16.1 |
| 260.0661 | 6.0 | | 116.0489 | 13.5 |
| 71.0487 | 4.5 | | 118.0644 | 12.6 |
| 99.0436 | 4.1 | | 170.0707 | 11.1 |
| 61.0280 | 4.0 | | 143.0597 | 10.3 |
| 213.0891 | 3.3 | | 134.0595 | 8.3 |
| 75.0436 | 3.2 | | 198.0655 | 6.0 |
| 160.0864 | 3.0 | | 187.0501 | 4.7 |
| 73.0280 | 2.8 | | 145.0754 | 4.2 |
| 206.0919 | 2.4 | | 131.0598 | 3.8 |
| 84.0803 | 2.2 | | 216.0767 | 3.4 |
| 230.0920 | 2.1 | | 106.0647 | 3.3 |
| 186.0656 | 2.1 | | 171.0548 | 3.1 |
| 53.0383 | 2.1 | | 117.0568 | 2.8 |
| | | | 119.0489 | 2.8 |

Table S4. Product ions and relative abundances of the MS$^2$ fragmentation of m/z 397.1358 at HCD 55 and MS$^2$ (397.1358) HCD 55 followed by MS$^3$ (263.0779) HCD 55.

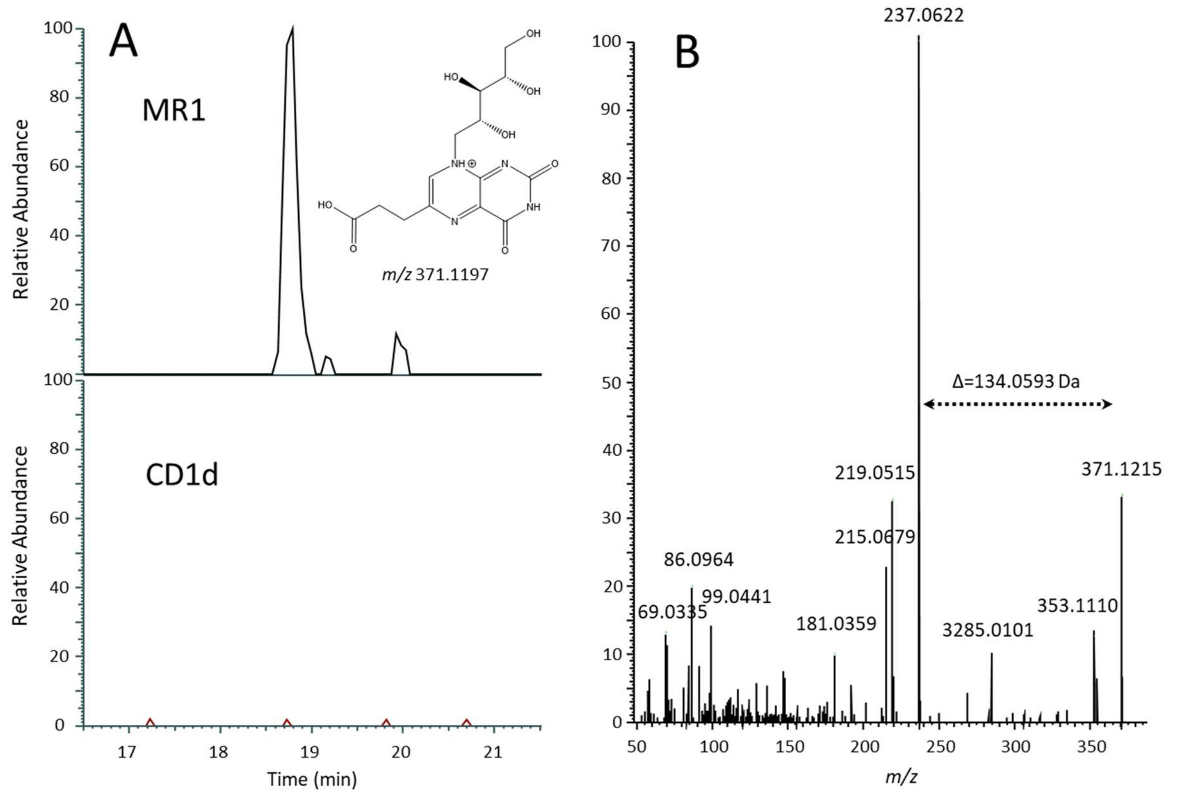


Figure S4. MS detection of a 370.1124 Da analyte. Panel A is the *m/z* 371.1197 extracted ion chromatogram with 5 ppm mass accuracy of the MR1 sample (top) and the CD1d sample (bottom). Panel B is the DDA $MS^2$ spectrum acquired at retention time 18.80 minutes of the MR1 sample with a prominent peak at *m/z* 237.0622 consistent with the neutral loss of ribityl.

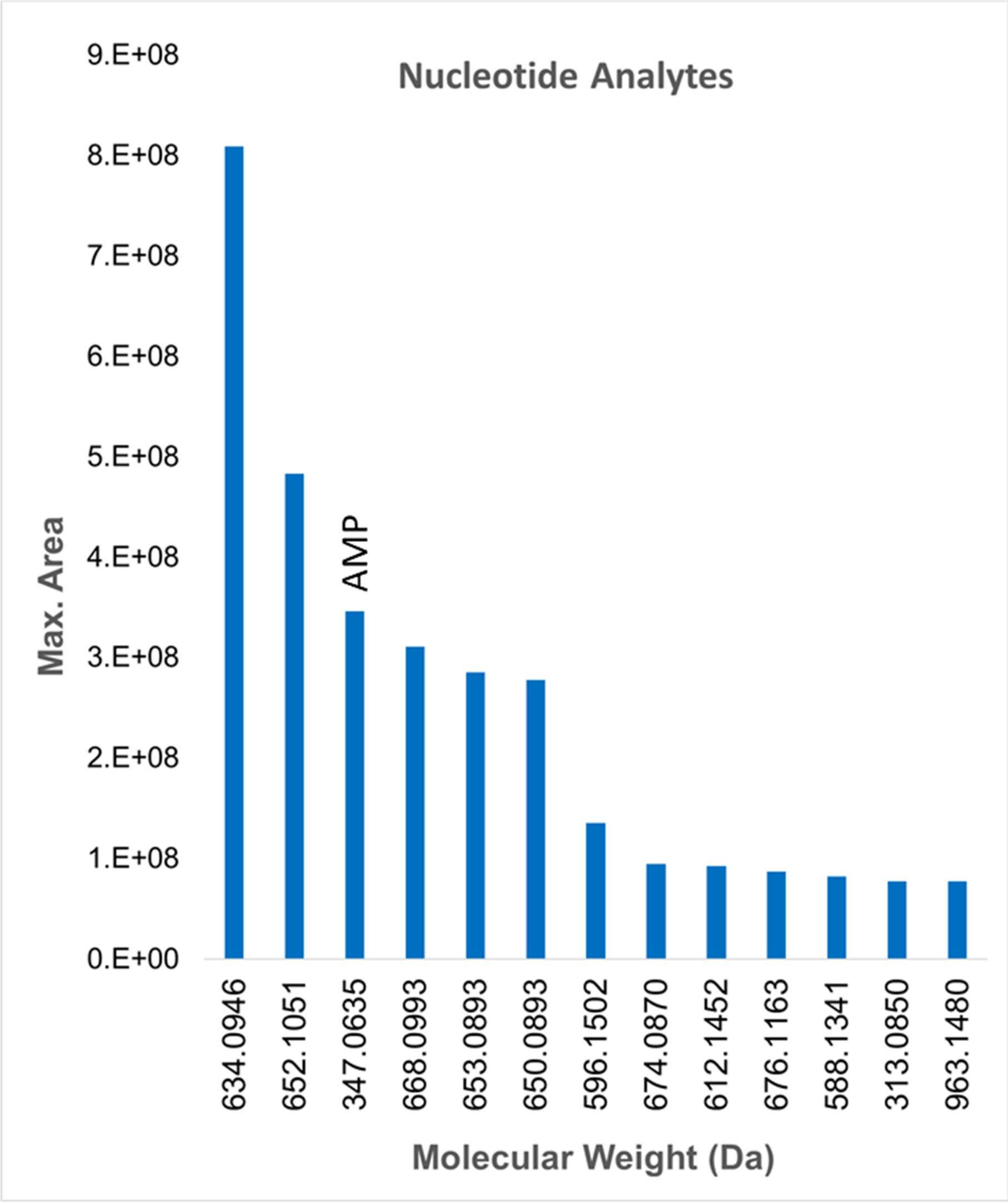


Figure S5. Extracted protonated ion chromatographic peak areas for most abundant nucleobase phosphate analytes listed in Table 3. AMP is adenosine monophosphate.

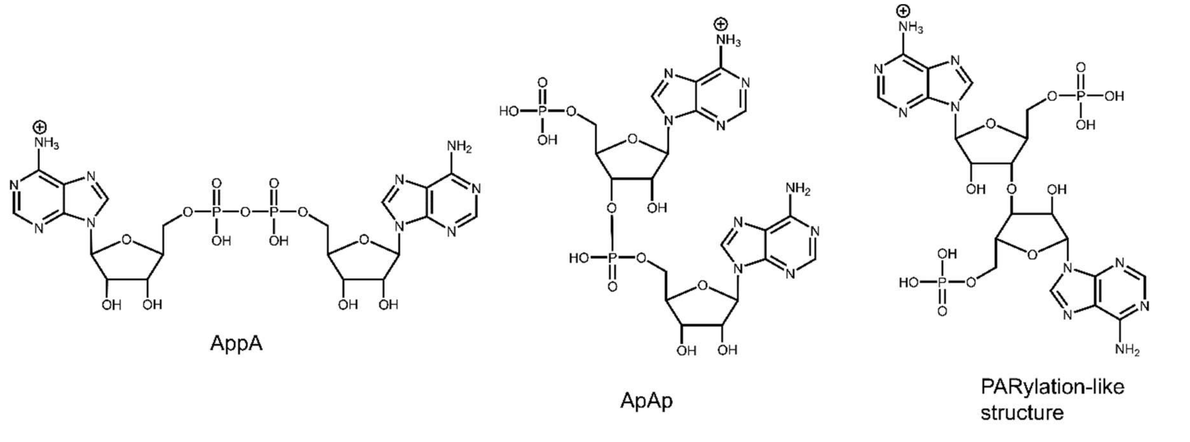


Figure S6. Possible structures for the assignment of the 676.1163 Da analyte.

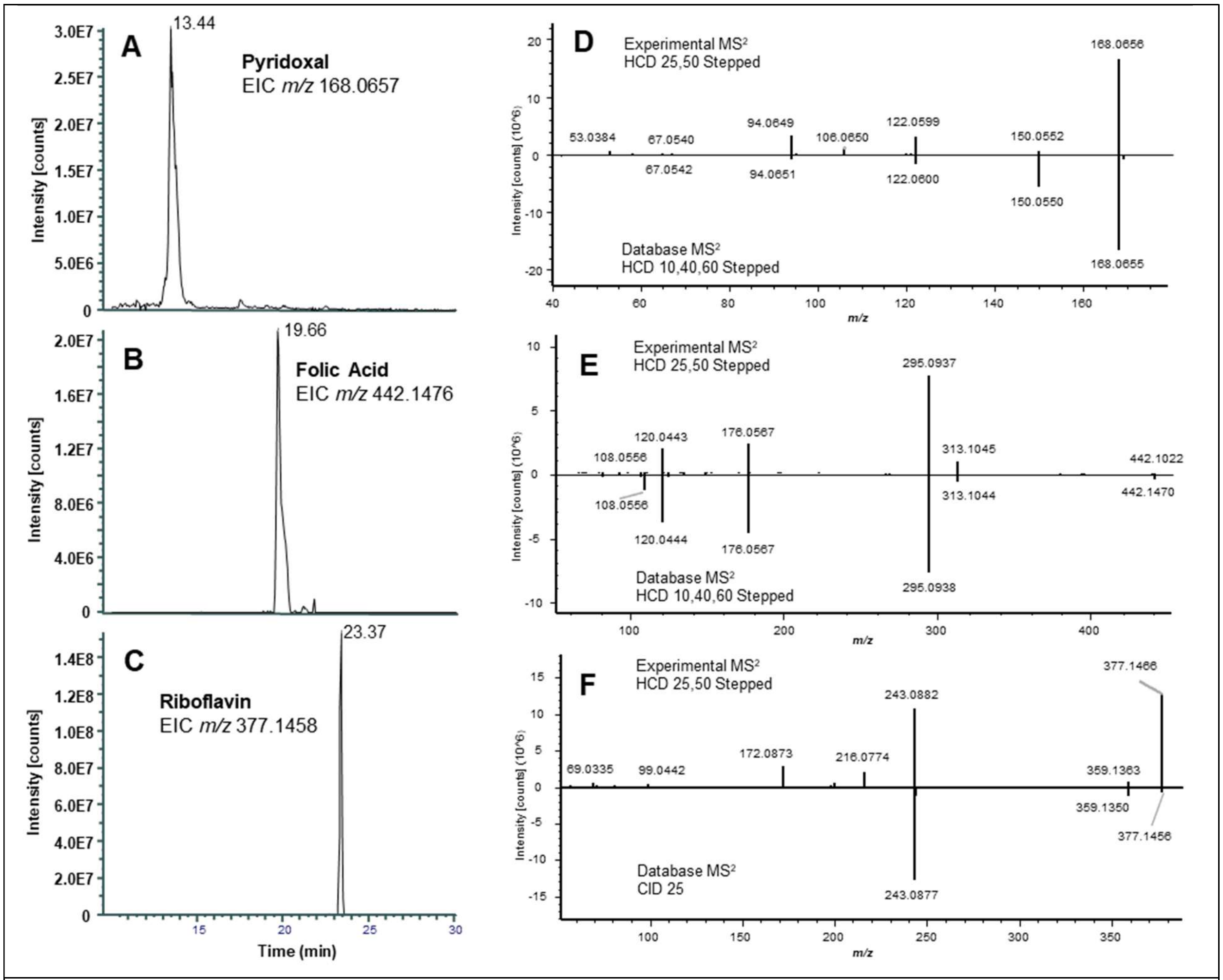


Figure S7. Manual confirmation of the Compound Discoverer identification of pyridoxal, folic acid and riboflavin in sMR1 samples. Panel A, B and C are the extraction ion chromatograms with 5ppm mass tolerance of pyridoxal (*m/z* 168.0657), folic acid (*m/z* 442.1476) and riboflavin (*m/z* 377.1458), respectively. Panels D, E and F are the experimental spectrum (top) and the corresponding spectrum from the mzCloud database (bottom) for pyridoxal, folic acid, and riboflavin, respectively.
